# Sequence space of Xpo1-dependent NESs reveals a functional affinity ceiling

**DOI:** 10.64898/2026.07.31.742000

**Authors:** Oleh Rymarenko, Trevor Huyton, Dirk Görlich

## Abstract

Exportin 1 (Xpo1/Crm1) exports hundreds of proteins from the nucleus to the cytoplasm. It recognizes nuclear export signals (NESs) with 4-5 hydrophobic Φ residues separated by spacer residues. Here we explored the sequence space of the most common NES class by a high-throughput ratiometric scoring of Xpo1-binding, combining phage display with deep sequencing. This provided a positional preference map and revealed that not only the Φ-positions but also spacer and flanking residues are critical for NES activity. We validated these data *in vivo* and with a new equilibrium affinity measurement that exploits the competition for Xpo1 when NES·Xpo1·RanGTP complexes partition into an FG phase. Guided by these preferences, we designed peptides that satisfy the established NES consensus but fail to confer export. Conversely, we engineered NESs that bind Xpo1 with low picomolar affinity — explained by a crystal structure. Such extreme binders, however, are no longer released from Xpo1 and block export *in trans*, explaining why natural NESs remain modest in affinity. Our data provide a framework for predicting, identifying, and engineering NESs and other peptide-based signals.

## Introduction

The nuclear envelope (NE) divides eukaryotic cells into two main compartments – the nucleus and the cytoplasm. The cell nucleus harbors the genome and hosts the processes associated with its maintenance, as well as the initial steps of gene expression. The cytoplasm, on the other hand, is specialized in protein synthesis. This makes a controlled exchange of macromolecules between the two compartments essential. The exchange occurs through the nuclear pore complexes (NPCs) that span the NE and feature a 8-fold rotational symmetry (Feldherr, 1962; Stevens and Swift, 1966; Gall, 1967; for latest structural models see Schuller *et al*., 2021; Bley *et al*., 2022; Petrovic *et al*., 2022; Zhu *et al*., 2022; Obarska-Kosinska *et al*., 2026). NPCs are composed of so-called nucleoporins (reviewed by Knockenhauer and Schwartz, 2016; Hampoelz *et al*., 2019; Mobbs *et al*., 2026). These include the FG-nucleoporins with long intrinsically disordered FG domains containing numerous FG (phenylalanine-glycine) dipeptide motifs (Hurt, 1988; Wente *et al*., 1992; Powers *et al*., 1997).

Active nucleocytoplasmic transport is typically mediated by the interplay of three components: (i) the FG phase, an FG domain condensate that serves as the NPC permeability barrier (Ribbeck and Görlich, 2002; Frey and Görlich, 2007; Schmidt and Görlich, 2015), (ii) the RanGTPase system that provides energy input and confers transport directionality (Bischoff and Ponstingl, 1991; Görlich *et al*., 1996; Weis *et al*., 1996; Izaurralde *et al*., 1997), and (iii) shuttling nuclear transport receptors (NTRs) that bind cargos in a RanGTP-controlled fashion and confer facilitated translocation through the FG phase (reviewed by Schmidt and Görlich, 2016; Mobbs *et al*., 2026).

NTRs include importins that import cargo into the nucleus as well as exportins that mediate transport in the opposite direction (reviewed by Güttler and Görlich, 2011; Kimura and Imamoto, 2014; Matsuura, 2016; Kehlenbach and Chook, 2025). Exportins bind cargos inside nuclei cooperatively with RanGTP (Fornerod *et al*., 1997; Kutay *et al*., 1997). In the case of exportin 1 (Xpo1, also known as Crm1), cargo affinity is enhanced by about 1000-fold by nuclear RanGTP and *vice versa* (Paraskeva *et al*., 1999; Güttler *et al*., 2010). Hydrolysis of the Ran-bound GTP in the cytoplasm switches Xpo1 to a low-affinity conformation (Monecke *et al*., 2013), allowing cargo release and thus directional transport.

Since exportin-bound RanGTP is initially resistant to RanGAP, GTPase activation requires the co-activators RanBP1 or RanBP2 (Bischoff *et al*., 1995; Bischoff and Görlich, 1997; Kutay *et al*., 1997; Paraskeva *et al*., 1999). These co-activators bind RanGTP within the complex, weaken the NTR–RanGTP interaction, and ‘present’ a transiently released RanGTP·RanBP1 (or RanBP2) subcomplex to RanGAP. Only then can GTP hydrolysis be triggered, rendering export-complex disassembly irreversible. Structures of RanGTP bound to Xpo1 (Monecke *et al*., 2009), RanGAP (Seewald *et al*., 2002), and the RanBD4 of RanBP2 (Vetter *et al*., 1999) explain why disassembly has to occur in this sequence. At least in yeast, RanBP1 might trigger cargo release from Xpo1 even before GTP hydrolysis locks in the disassembled state (Koyama and Matsuura, 2010).

Xpo1 exports ∼1000 different proteins from mammalian cell nuclei (Thakar *et al*., 2013; Kirli *et al*., 2015; Mackmull *et al*., 2017), including numerous regulatory factors. It further exports pre-40S and pre-60S ribosomal particles (Ho *et al*., 2000; Gadal *et al*., 2001; Thomas and Kutay, 2003) as well as certain viral genomes (e.g., HIV and influenza; see Fischer *et al*., 1995; Wolff *et al*., 1997; Elton *et al*., 2001). Moreover, Xpo1 returns many cytoplasmic components, such as translation factors, RanBP1, and RanGAP, back to the cytoplasm if they have leaked into the nuclei (Feng *et al*., 1999; Künzler *et al*., 2000; Bohnsack *et al*., 2002; Kirli *et al*., 2015).

Xpo1 is the best studied nuclear transport receptor thanks to genetic studies (Stade *et al*., 1997) and the early discovery of its specific inhibitor leptomycin B or LMB for short (Hamamoto *et al*., 1983; Nishi *et al*., 1994; Fornerod *et al*., 1997; Fukuda *et al*., 1997; Ossareh-Nazari *et al*., 1997; Wolff *et al*., 1997). LMB is cell-permeable and can therefore block Xpo1 in cell culture experiments. It covalently modifies cysteine 529 of *S. pombe* Xpo1 (Kudo *et al*., 1999), which corresponds to C528 of the human protein and resides directly in the NES-binding site (Dong *et al*., 2009; Monecke *et al*., 2009; Sun *et al*., 2013). This cysteine corresponds to a threonine (T539) in *S. cerevisiae* Xpo1, and this exchange explains the LMB-resistance of this yeast (Neville and Rosbash, 1999).

The study of Xpo1 was also aided by the fact that it recognizes rather simple (leucine-rich) nuclear export signals, called NESs for short (Wen *et al*., 1994; Fischer *et al*., 1995). NESs are located in disordered protein regions and are therefore transplantable. They comprise a crucial set of hydrophobic amino acid residues (called Φ residues) separated by variable spacer residues. Prototypes are the NES of the protein kinase A inhibitor PKI (PKI NES; Wen *et al*., 1994) with four essential hydrophobic amino acids (Φ^1^-Φ^4^) and the NES of the HIV-1 Rev protein (Rev NES; Fischer *et al*., 1995). As more NESs have been discovered and characterized, additional sequence motifs have been described for them (Kosugi *et al*., 2008).

The first crystal structures of Xpo1, with snurportin 1 (SPN1) bound as a cargo, revealed that the NES-binding site of Xpo1 comprises not just four, but five pockets for hydrophobic Φ residues (Dong *et al*., 2009; Monecke *et al*., 2009). Likewise, the fifth hydrophobic Φ-residue (Φ^0^) has been shown to increase the Xpo1 affinity of the PKI NES, and the shorter Rev NES also docks with five Φ-residues to the exportin (Güttler *et al*., 2010). Indeed, the same five pockets can accommodate NESs with different Φ-spacings. However, this requires that they bind in different conformations, namely with an α-helical (PKI NES Φ^0^L) or an extended (Rev NES) N-terminus (Güttler *et al*., 2010). Yet another NES class binds in reverse orientation (Fung *et al*., 2015).

The PKI NES class has been described by a Φ^0^xxΦ^1^xxxΦ^2^xxΦ^3^xΦ^4^ consensus, where ‘x’ denotes variable spacer amino acids. The highest-affinity NESs reported to date (the so-called supraphysiological or superNESs) all follow this pattern, although the early studies missed the relevance of the affinity-enhancing Φ^0^ residue (see Engelsma *et al*., 2004, 2008; Güttler *et al*., 2010; Fung *et al*., 2015). This may suggest that the PKI class represents the energetically most favorable spacing pattern. It should therefore allow for the greatest sequence diversity, which in turn is consistent with the majority of NESs discovered so far belonging to the PKI class (Lee *et al*., 2019).

The highly redundant PKI NES class consensus alone covers an astronomical number (∼10^13^) of different sequences that may or may not represent genuine Xpo1-binders. It is indeed an intriguing question which additional features enhance or impede Xpo1-binding and NES function, especially considering that most of the potential variability originates from the poorly studied inter-Φ spacers.

To obtain a comprehensive overview of the sequence space of the PKI NES class, we set up phage display as a quantitative, high-throughput, ratiometric scoring of Xpo1 binding. This involved NES library construction, phage selection on immobilized Xpo1, highly specific protease elution, and deep sequencing to quantify all NES species in the starting and bound material. We applied this approach to a library of all possible single-point mutations of the PKI NES. This confirmed and quantified the known Xpo1 preferences at the Φ positions and, in addition, revealed preferences for all flanking and spacer residues, showing that their impact goes far beyond what was previously recognized. We validated these findings not only by testing export activity *in vivo*, but also with a new equilibrium affinity measurement exploiting the competition for Xpo1 during FG-phase partitioning of NES·Xpo1·RanGTP complexes. Using the discovered preferences, we rationally designed NES-like sequences that match the established consensus but do not function as effective NESs. Conversely, we designed new sequences that match the NES activity of the PKI NES while sharing no sequence identity with the prototype. Furthermore, we engineered sequences that bind Xpo1 with extreme affinity, far exceeding all previously reported superNESs. We discuss why such binders could hardly be obtained by direct selection. We also provide a crystal structure that confirms the expected binding mode and explains this exceptionally strong interaction. Finally, we document the essentially irreversible Xpo1 binding of such peptides *in vivo*.

## Results

### Sequence space of PKI-type NESs

To elucidate the sequence space of the PKI NES class by mutational analysis and high-throughput ratiometric affinity measurement, we combined phage display (Smith, 1985; Bass *et al*., 1990) with Xpo1 affinity selection (Fig. 1A). An M13 phagemid was constructed to display a pIII-fused PKI NES library, including flanking residues and all possible single point mutations of this 15-residue long sequence (Fig. 1B-C). In this approach, each NES peptide gets physically linked to its encoding DNA that is packaged within the pIII-containing phage. pIII is a pentameric protein, and its multivalency complicates a quantitative readout. To minimize avidity effects, we “diluted” the peptide-fused pIII molecules with non-fused ones from the helper phage (Vieira and Messing, 1987) thus keeping the display levels on the pIII pentamer low. Since such dilution is random, the initial phage pool also included phages without any displayed peptide (and containing only non-fused pIII). Furthermore, display levels are likely to vary among different peptides due to differences in the efficiency of translation and secretion, or possibly even proteolysis of the NES fusion. This makes it difficult to distinguish between an inefficient display of a given peptide and inefficient binding to Xpo1. In other words, non-displaying phages introduce a bias, precluding accurate ratiometric comparisons between different NESs.

**Figure 1.**
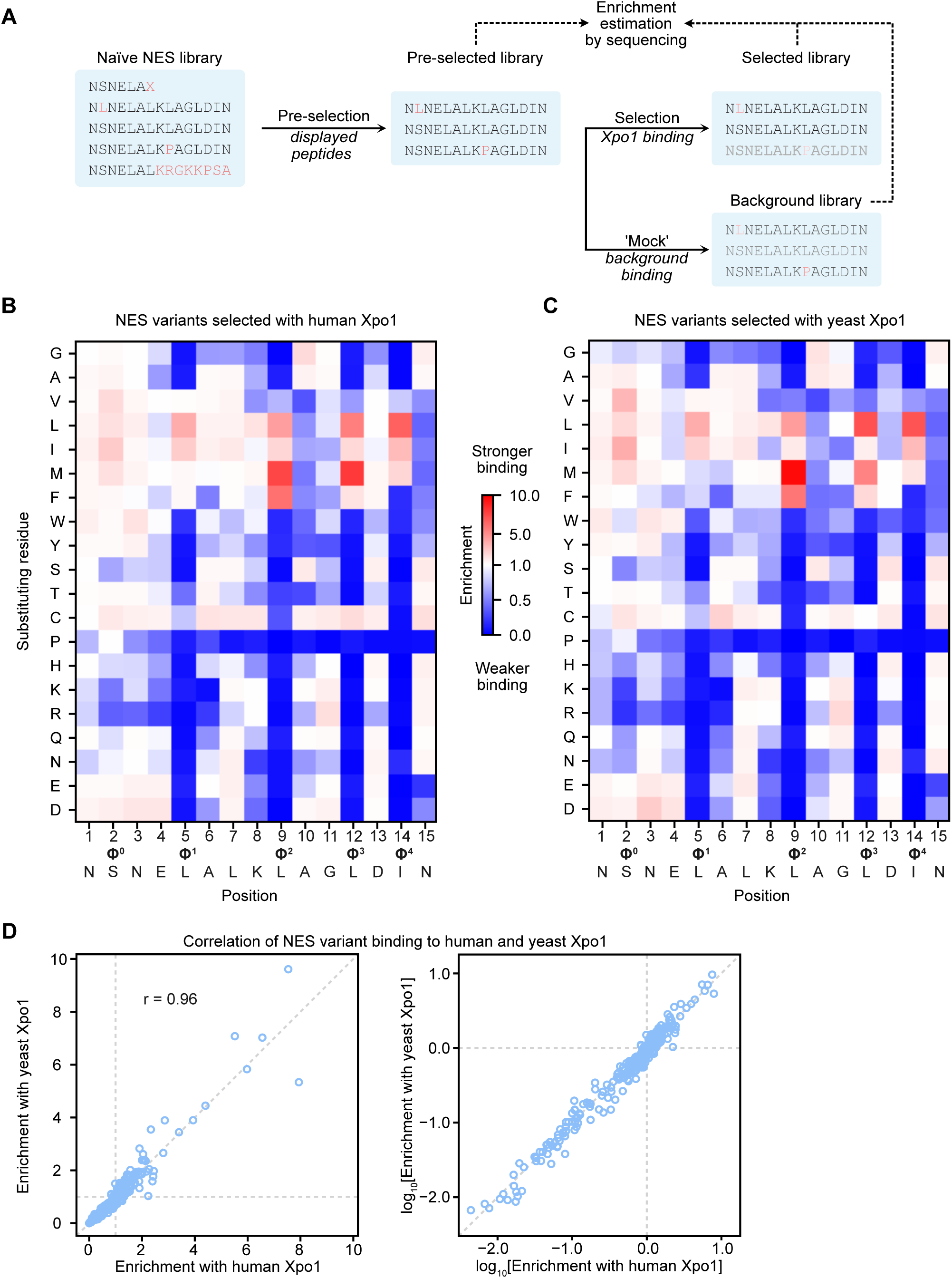
Residue preferences of human and yeast Xpo1 across PKI-type NESs. (**A**) Workflow for mutational scanning across the PKI NES sequence, phage-display selection of Xpo1-binding variants, and quantification of binding by deep sequencing. (**B**, **C**) Heatmaps showing the positional preferences of human Xpo1 (B) or S. cerevisiae Xpo1 (C) for binding a PKI-type NES. At a given NES position, red indicates residues that confer stronger exportin binding; blue indicates weaker binding. See main text and Methods for a full description; numerical values are provided as Appendix Tables 1 and 2. (**D**) Correlation of human and yeast Xpo1 positional preferences, shown in linear (left) and log-log (right) representations. The 300 data points comprise all single-point mutations depicted above, with the x- and y-coordinates giving the human and yeast Xpo1 preference values, respectively.

We overcame this problem by pre-purifying peptide-displaying phages utilizing an affinity capture and proteolytic release step, using a SUMO^Eu^ cleavage module (Vera Rodriguez *et al*., 2019) fused upstream of the NES library. These pre-selected phages were then allowed to bind (in the presence of RanGTP) to human or *S. cerevisiae* Xpo1, which in turn had been immobilized on magnetic beads via an xLC3B cleavage module. Phages were then eluted by releasing Xpo1 from the beads, using the xAtg4 protease for xLC3B cleavage (Frey and Görlich, 2015) To account for any residual background phages, a selection without Xpo1 (but with the identical affinity tag containing the xLC3B cleavage module immobilized on the beads) was performed in parallel. Quantification of the eluted phages showed, however, that this background was very low – only 1/3000 of the samples with Xpo1.

The selected libraries, as well as the pre-selected naïve library, were subjected to Illumina sequencing (Bentley *et al*., 2008) of the NES-encoding phagemid inserts. The abundance of a given NES sequence in each library was estimated from its read counts. Normalized enrichments in the Xpo1-bound fractions were then calculated for each NES variant (see Materials and methods for details) and used for the analysis (Fig. 1B and C, and below). This method does not provide absolute Xpo1 affinities but is ideal for a ratiometric ranking of binding strength: all numbers derive from the same binding reaction, avoiding the variability of parallel or sequential assays.

### Details of Xpo1 preferences

The starting point for library construction was the wild-type PKI NES (Wen *et al*., 1994), a medium-strength Xpo1 binder (Güttler *et al*., 2010; Fung *et al*., 2015) allowing the analysis to reveal both deleterious mutations and mutations that enhance the interaction with the exportin. The strongest positive selection was evident at and around Φ^0^ (Fig. EV1A). This is consistent with the fact that a suboptimal serine, and not a hydrophobic residue, occupies this position in the wild-type PKI NES (Güttler *et al*., 2010). All other Φ positions showed much less selective pressure for an exchange, indicating that they were already occupied by favored residues.

The heatmaps of the Fig. 1B-C show the enrichment values (available in tabular form as Dataset 1B and Dataset 1C) for all mutations. Thus, they represent a compact summary of amino acid preferences for all positions of the NES sequence. For example, any exchange of a hydrophobic Φ^1^-Φ^4^ residue for a hydrophilic one is deleterious. Yet, these positions differ in their preferences. For example, methionine or phenylalanine are preferred at Φ^2^, whereas leucine is optimal at Φ^1^ and Φ^4^. This is in excellent agreement with previous findings (Kosugi *et al*., 2008; Güttler *et al*., 2010) and validates our novel approach.

As mentioned above, the wild-type PKI NES has a serine at position 2 (Φ^0^) instead of a hydrophobic residue. As expected, we observed a strong selection for hydrophobic residues (I, V, M, L, C) at this position. A similar, but milder, hydrophobic selection is evident at positions 1 and 3, in particular for tryptophan (N1W and N3W). Tryptophan is favored only at these positions, and we assume that its sidechain docks into the Φ^0^ pocket, changing the characteristic PKI NES Φ^0^xxΦ^1^ pattern to a Φ^0^xxxΦ^1^ or a Φ^0^xΦ^1^ spacing and altering the conformation of the bound peptide. Negatively charged residues are also favored in positions 1 to 4, probably due to charge complementarity to adjacent lysines and histidines in Xpo1 (Güttler *et al*., 2010). We see this as a further validation of our strategy.

Proline is tolerated only at the (non-essential) Φ^0^ position, where it could provide a similar hydrophobic contact as a Φ-residue. It is deleterious at all other Φ and spacer positions and even at position 15 after Φ^4^, consistent with the previously reported observations (Kosugi *et al*., 2008; Güttler *et al*., 2010). A likely explanation is incompatibility with the secondary structure of the bound NES peptide. The spacer positions showed a number of striking additional constraints. For example, glycine is disfavored in the Φ^1^–Φ^2^ and Φ^3^–Φ^4^ spacers, perhaps because of a destabilizing effect on the secondary structure. Interestingly, hydrophobic residues are selected against in the Φ^2^–Φ^3^ spacer and at position 15, where acidic residues are also strongly disfavored.

Basic residues, on the other hand, are disfavored in the N-terminal half of the NES. Cysteine appears to be preferred at all spacer positions and even at Φ^3^. Leucine and methionine are selected for at spacer position 8, and arginine and lysine at position 11. Conversely, threonine is disfavored at position 10. Overall, the contributions and constraints of each residue at each position of a PKI-type NES to Xpo1 binding have now been assessed and quantified.

### Extreme evolutionary conservation of positional NES preferences

Having performed the experiment using Xpo1 from humans and *S. cerevisiae* as baits, we were able to evaluate the evolutionary conservation of the Xpo1 preferences. The heatmap patterns from the Fig. 1B and 1C look strikingly similar, and the residue-specific preferences are highly correlated (r = 0.96, Fig. 1D). The most prominent difference is seen at Φ^3^, (Fig. 1B–C, and EV1B) where human Xpo1 prefers methionine over leucine, whereas yeast Xpo1 has the opposite preference relationship. This difference appears to be related to their different leptomycin B sensitivities. Φ^3^ contacts the LMB-reactive cysteine 528 of human Xpo1, which is substituted by threonine 539 in the yeast exportin (Fig. EV1C, D). This excellent cross-validation between the human and yeast datasets demonstrates that our experimental approach reliably detects even small structural details. The idea of general preference conservation is further supported by the fact that PKI Φ^0^L NES binds with virtually identical affinities to both human and yeast Xpo1 (Fig. EV1E). However, it cannot be excluded, that the proteins from the two species have differing affinities for other NESs or the different ranges of affinities altogether.

Although impressive, such strong evolutionary conservation of Xpo1 preferences is not unexpected. Xpo1 is an essential protein (Adachi, 1989; Stade *et al*., 1997) and is highly conserved across the eukaryotes. The sophisticated allosteric coupling between the binding of RanGTP and cargo (Monecke *et al*., 2009, 2013; Saito and Matsuura, 2013) leaves little room for sequence variability at the NES-binding pocket. Furthermore, since hundreds of proteins rely on Xpo1 for nuclear export (Thakar *et al*., 2013; Kirli *et al*., 2015; Mackmull *et al*., 2017), a change in Xpo1 preferences would require a concomitant change in the NESs of these proteins, which is highly unlikely to occur in the normal course of incremental evolution.

### An FG phase-based, ratiometric assay for Xpo1-affinities of NESs

We have so far identified NES spacer mutations that increase or decrease Xpo1 binding, with phage display providing a robust ranking of binding strength. Because this ranking involved affinity chromatography with extensive washing, the scores are biased toward an off-rate selection. Obtaining true affinities therefore requires either a method that determines both on- and off-rates, or one that operates at equilibrium. However, we found the standard affinity-measurement methods were problematic for this task. For example, the extreme cooperativity between NES and RanGTP binding makes it impractical to derive K_D_ values for the Xpo1– NES interaction from kinetic biolayer interferometry or surface plasmon resonance measurements. Fluorescence anisotropy (Dandliker and Feigen, 1961), previously used for NES affinity measurements (Güttler *et al*., 2010; Fung *et al*., 2015; Fung and Chook, 2022), is reliable only over a limited affinity range, as we show below for high-affinity NESs.

To measure a wide range of NES affinities in a functional context, we therefore established a new assay based on the FG phase (Ribbeck and Görlich, 2002; Frey and Görlich, 2007; Frey *et al*., 2018). While inert macromolecules on their own remain phase-excluded, they become highly soluble in this phase when captured by a cognate NTR. This phenomenon can be recapitulated using ‘FG particles’ assembled from a phase-separating Nup98 FG domain (Schmidt and Görlich, 2015).

Indeed, a PKI NES-EGFP fusion stays excluded from the phase, yielding a partition coefficient of ∼0.2. In the presence of Xpo1 and RanGTP, however, it accumulates to a partition coefficient of >1000 (Fig. 2A; Ng *et al*., 2021). For our new assay, we let two NESs (fused to distinct fluorescent proteins) compete for a limiting concentration of Xpo1, let their Xpo1-bound fractions partition in the FG phase, and measure the accumulation at equilibrium by confocal laser scanning microscopy (Fig. 2B–C). The ratio of the Xpo1 affinities of the two NESs can then be read directly from the ratio of their FG phase:buffer partition coefficients (for derivation see Appendix Supplementary Materials).

**Figure 2.**
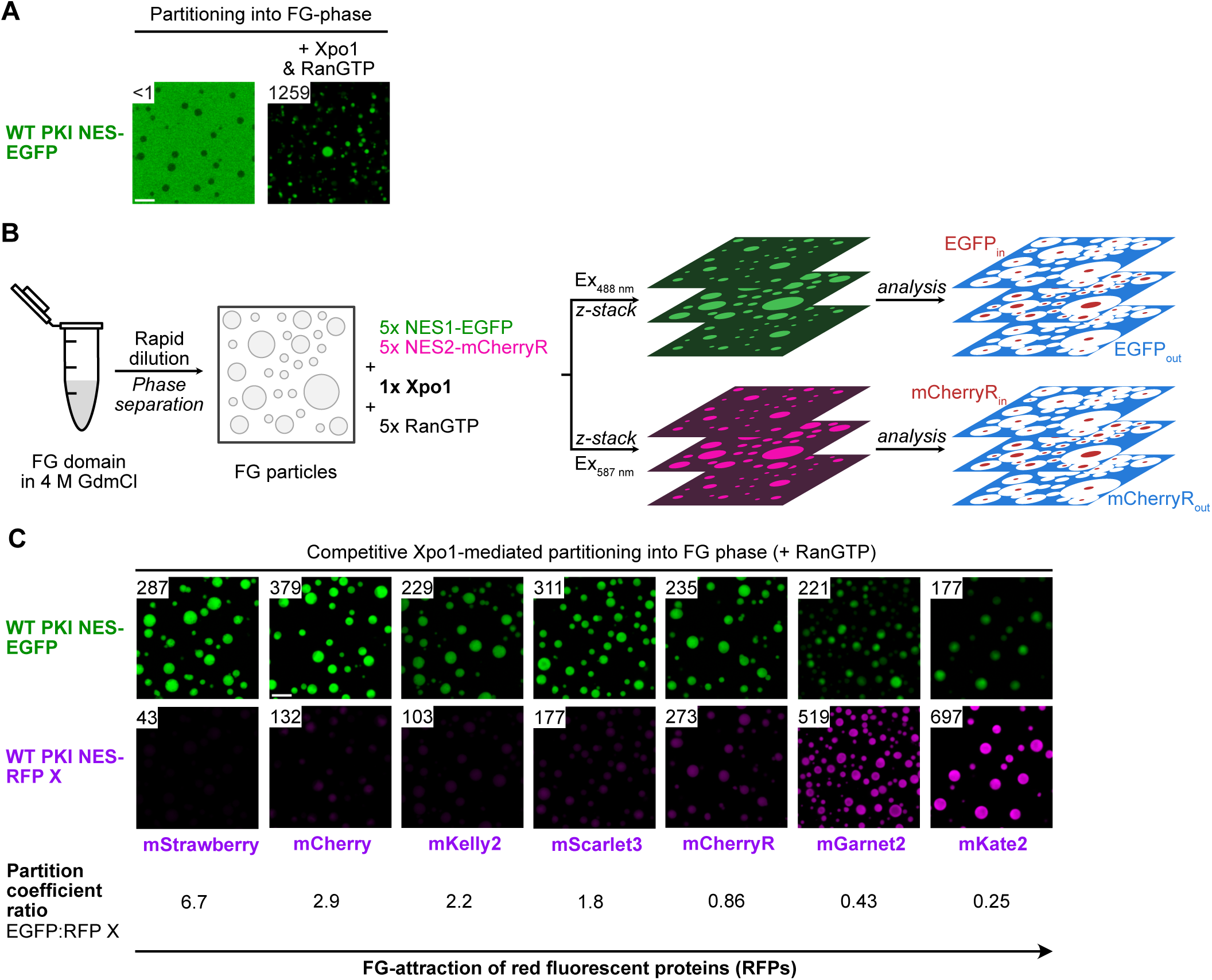
Relative NES affinities measured by competition for Xpo1 during FG phase partitioning. (**A**) Partitioning of wild type (WT) PKI NES-EGFP into a Nup98 FG phase, in the presence or absence of human Xpo1 (equimolar to the EGFP-NES fusion) and excess hRanGTP(5–180, Q69L). The partition coefficient (the ratio of the mean signal inside the particles to that in the adjacent buffer) is shown in the top-left corner of each image (here <1 without Xpo1, 1259 with Xpo1). Details of the procedure and quantification are given in the Methods. Scale bar, 10 µm. (**B**) Workflow of the competitive assay. Two NESs, fused to EGFP and to mCherryR, are pre-mixed, then supplemented with a limiting amount of Xpo1 and subsequently with an excess of RanGTP; FG particles, prepared by rapid dilution of a concentrated FG-domain stock from denaturant (4 M guanidine hydrochloride, GdnHCl), are added last. After the partition equilibrium is reached, the particles are imaged as z-stacks and partition coefficients computed. Because Xpo1 is limiting, it carries the NES-fusions in proportion to their affinities into the phase; the ratio of the two partition coefficients therefore reports their relative Xpo1 affinity. (**C**) The same NES (WT PKI), fused either to EGFP or to a red fluorescent protein, competed for Xpo1. Because the NESs are identical, the two fusions should partition equally (ratio 1); any deviation reflects a partitioning bias from the fluorescent fusion partner. The ratio deviates strongly from 1 for all red fluorescent proteins except mCherryR, engineered here as a close match to EGFP. The ratios place the red fluorescent proteins on a simplified FG-attractiveness scale below the panel. Scale bar, 10 µm.

For a wide dynamic range, the assay requires clean spectral separation of the two fluorescent proteins (FPs). We therefore chose EGFP and a red fluorescent protein (RFP) as our reporter pair. The assay is also most straightforward to interpret if the two different fluorescent fusion partners do not bias partitioning in the FG phase. This detail matters. For example, mCherry is more strongly repelled from the phase than EGFP, reflecting their different surface amino acid compositions (Frey *et al*., 2018). Accordingly, Xpo1-mediated partitioning is 3-fold lower for a PKI NES fused to mCherry than to EGFP (Fig. 2C). Screening a panel of five alternative monomeric RFPs (Lambert, 2019) revealed no suitable match either: mGarnet2 and mKate2 were 2- to 4-fold more FG-attractive than EGFP, while mScarlet3, mKelly2, and mStrawberry were 2- to 7-fold more FG-repellent (Fig. 2C).

We therefore engineered a dedicated mCherry reference (mCherryR) to closely match the FG-partitioning potential of EGFP. This involved 17 (rather conservative) FG-attractive surface mutations (Frey *et al*., 2018), namely twelve K**→**R, three A**→**I, and two N**→**Q exchanges (see Appendix). Consequently, the ratio of the partitioning coefficients of EGFP and mCherryR fused to the same NES showed a narrow distribution centered at 0.85, independent of the Xpo1 affinity of the respective NES (Figs. 2C, 3B, EV2B–C; see also Fig. 8 below). As exemplified later, the assay provides robust ratiometric measurements of Xpo1-binding strength, resolving even subtle affinity differences. A single comparison covers up to four orders of magnitude — roughly two below and two above the reference. Chaining measurements through a panel of calibrated reference NESs extends the accessible range to six orders of magnitude (Fig. 8; Table 1).

**Figure 3.**
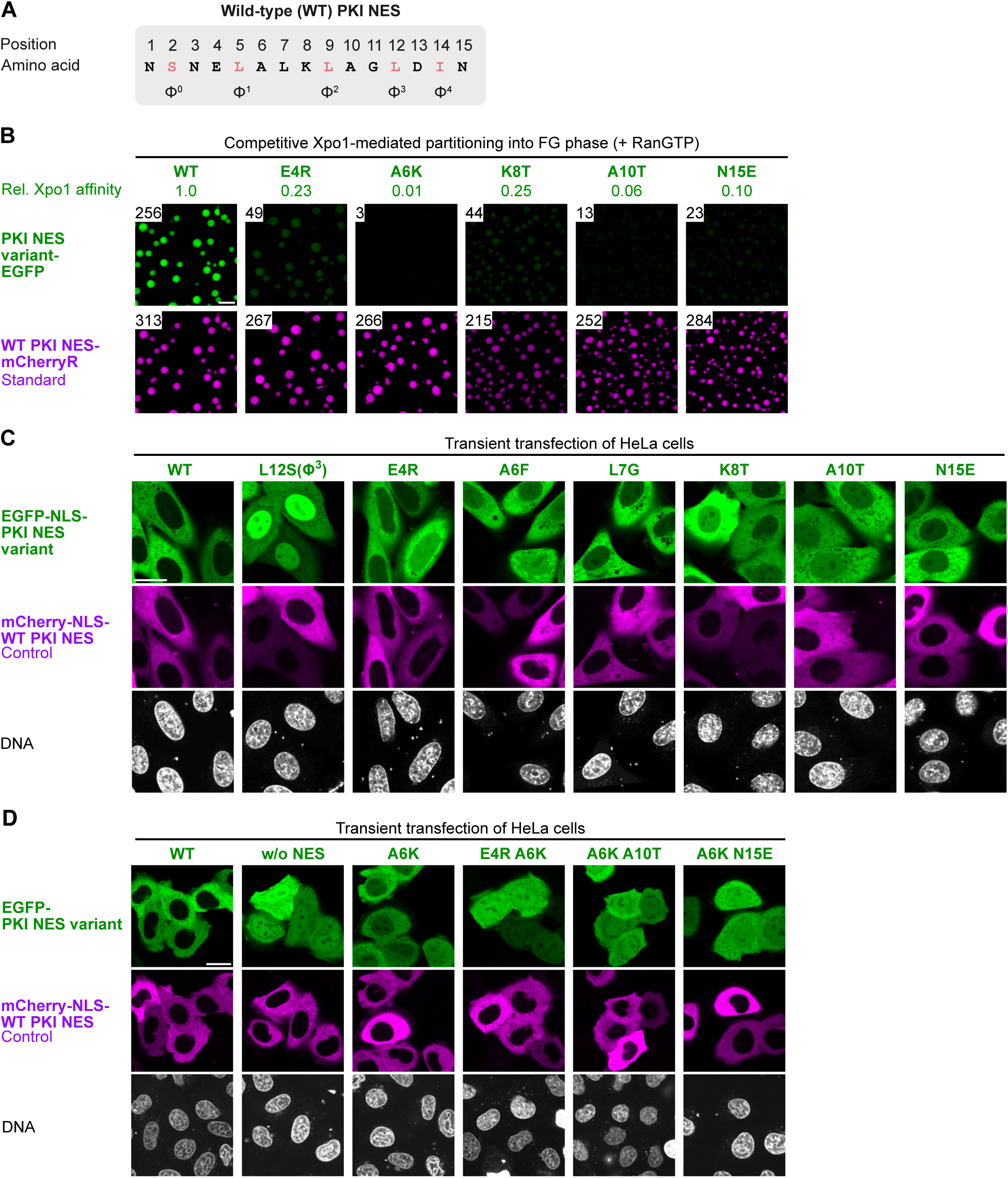
NES spacer mutations that weaken Xpo1 binding and export activity. Spacer mutations predicted from the preference map (Fig. 1B) to reduce Xpo1 binding indeed lowered Xpo1 affinity and impaired export activity in cells. Their combinations acted additively, abolishing export altogether. **(A)** Sequence of the PKI NES, given for orientation. **(B)** Analysis of EGFP-fused single spacer mutants followed the assay concept of Fig. 2. Each variant was allowed to compete with a WT PKI NES-mCherryR fusion for Xpo1-mediated FG phase partitioning. Partition-coefficient ratios (EGFP:mCherryR) were measured and converted into Xpo1 affinities (relative to the WT PKI NES) by dividing by the ratio of the included WT–WT pair (here 256:313 = 0.82). Scale bar, 10 µm. **(C)** Localization of single mutants in cells. Each was fused to the C-terminus of EGFP-SV40 NLS, expressed in HeLa cells, and imaged live by confocal microscopy. Efficient nuclear export is indicated by the reporter’s exclusion from the nucleus (marked by DNA stain). The mCherry-NLS-WT PKI NES control reports Xpo1-dependent export competence of the cells. Scale bar, 20 µm. **(D)** Localization of the A6K mutant and double mutants, fused to EGFP without an NLS, and compared the wildtype NES and a minus NES control. Without active import, localization reflects the balance of passive influx and NES-driven export, so only strong export impairment produces visible nuclear accumulation. Expressed and imaged as in (C). Scale bar, 20 µm.

**Table 1.**
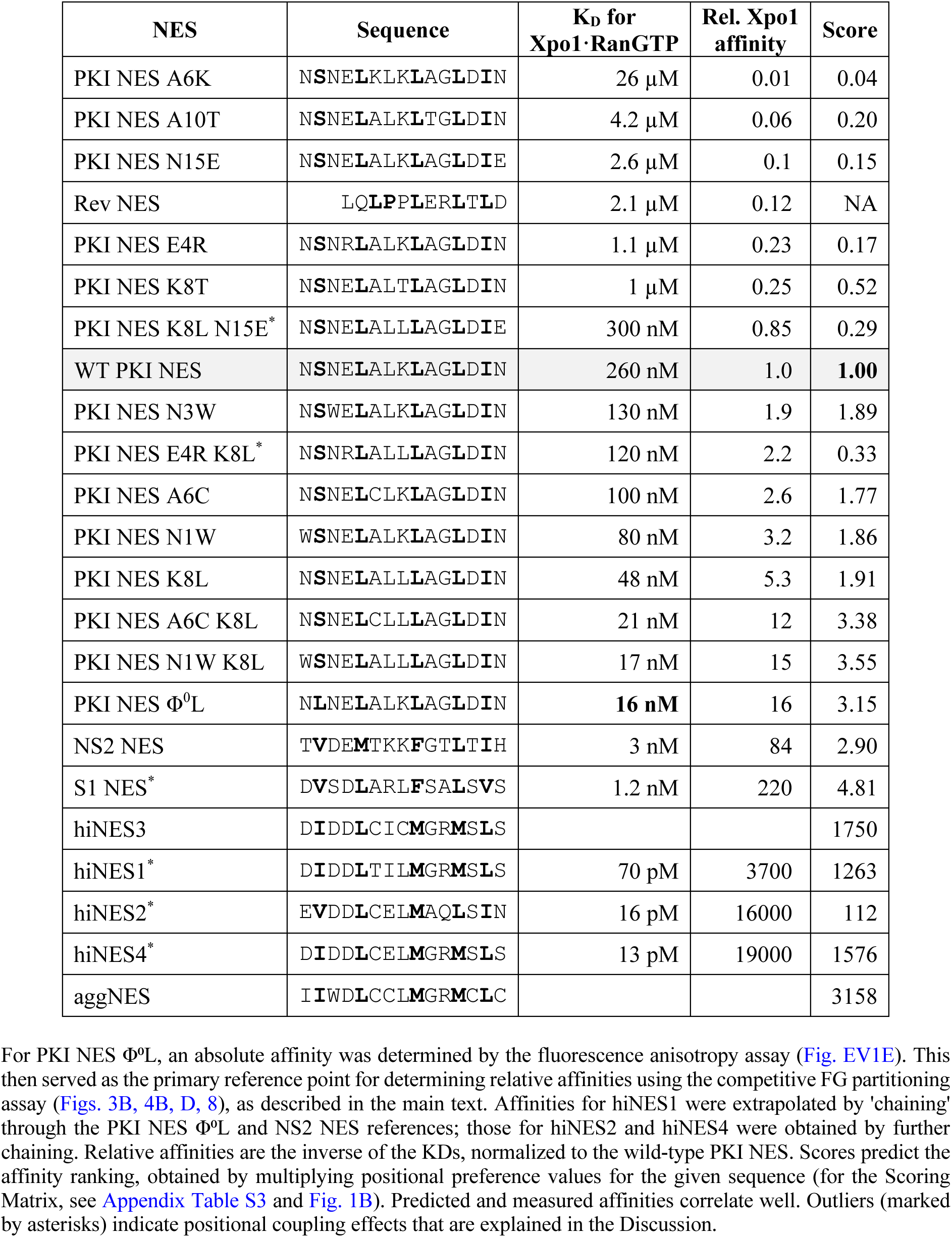
Affinities of selected NESs for the Xpo1·RanGTP complex.

We first applied the assay to putative negative spacer mutations and indeed observed 4- to 17-fold decreases in Xpo1 affinity for the E4R, K8T, A10T, and N15E exchanges (Fig. 3B, Table 1). The A6K mutation conferred an even more pronounced, ∼100-fold drop in affinity. Modeling onto Xpo1-bound PKI NES structures (Güttler *et al*., 2010; Koyama *et al*., 2014) revealed that a lysine at position 6 is poorly accommodated: it requires some structural rearrangement, remains highly rotamer-restricted even then, and its positively charged head group is electrostatically repelled by nearby exportin arginines.

### Disfavored spacer residues impair export activity *in vivo*

The preference data from our phage display-based approach along with the affinity measurements using the FG phase assay suggest a greater impact of the spacer residues on Xpo1 binding than previously acknowledged. To test how these effects play out *in vivo*, we transfected HeLa Kyoto cells with plasmids encoding EGFP fused to the SV40 NLS and a series of PKI NES variants (Fig. 3A, C). Since export must compete here against import through the SV40 NLS, this reporter assay is quite sensitive to deleterious NES mutations. As an internal control, cells were co-transfected with a construct encoding a fusion of mCherry with an SV40 NLS and the wild-type PKI NES, which normally localizes to the cytoplasm.

When spacer residues were mutated to strongly disfavored ones, the localization of a reporter construct was notably affected (Fig. 3C). Although not as dramatically as the highly detrimental LΦ^3^S mutation, which causes strong nuclear accumulation of the reporter, spacer mutations E4R, A6F, L7G, K8T, A10T, and N15E clearly reduce export efficiency and abolish the otherwise complete exclusion of reporter from the nucleus.

### Inter-Φ spacer mutations that enhance binding to Xpo1

Numerous PKI NES variants were positively selected during phage display with Xpo1 and RanGTP. Of these, we tested the spacer mutants N1W, N3W, A6C, and K8L in the FG phase assay (Fig. 4A, B). This revealed increased Xpo1 affinities of 3.2-fold (N1W), 1.9-fold (N3W), 2.6-fold (A6C), and up to 5.3-fold (K8L). Comparison of negative and positive mutations at the same positions (Fig. 3B and 4B) indicates an even larger 21-fold difference between the K8T and K8L, and a 260-fold between the A6K and A6C variants. For spacer positions previously denoted only as ‘x’ in the NES consensus, such large effects seem remarkable indeed.

**Figure 4.**
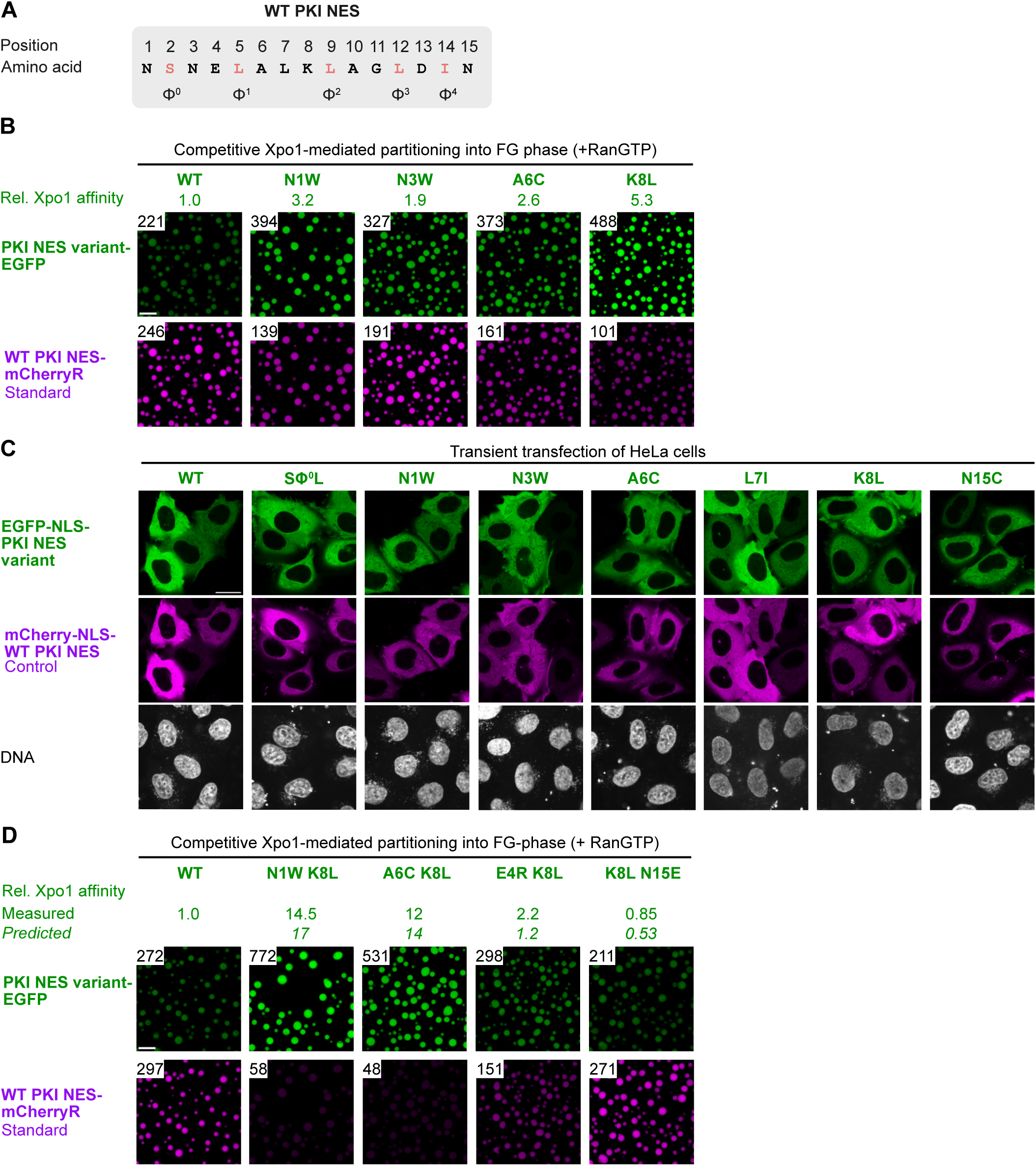
NES spacer mutations that strengthen Xpo1 binding. **(A)** PKI NES sequence. **(B)** Analysis of EGFP-fused single spacer mutants, as in Fig. 3B. Each exchange increased Xpo1 affinity, as predicted from the preference map (Fig 1B). **(C)** Localization of single mutants in cells, as in Fig. 3C. All variants exported efficiently — export is not limited by NES affinity in this range. **(D)** Analysis of double mutants, as in (B). Measured affinities are compared with those predicted from the single-mutation values (Figs. 3B, 4B). Combining two positive mutations was additive and brought the PKI NES to superNES level. A positive exchange (K8L) could also compensate a detrimental one (E4R or N15E). K8L rescued the E4R mutant (∼2-fold) beyond the predicted affinity, indicating a positional coupling.

We then tested these mutants in a transfection assay (Fig. 4A, C), together with the L7I and N15C spacer variants and the previously described affinity-enhancing SΦ^0^L mutant (Güttler *et al*., 2010). All variants conferred the same complete nuclear exclusion of the reporter as the wild-type PKI NES, confirming that the binding strength for Xpo1 is not limiting for export once it reaches the level of the PKI NES and that the tested mutations do not have significant detrimental effects on the PKI NES. This highlights the power of the competitive FG phase assay to distinguish affinity differences that cannot be discerned in the *in vivo* system.

### Combining spacer mutations of PKI NES

For NES predictions and peptide design, it is relevant whether mutations at different residues have predictable synergistic effects. This is expected if their ΔΔG contributions add up incrementally. In terms of affinities/K_D_ values, this equals a multiplication of mutation effect factors (according to Gibbs’ equation). In such a case, we should be able to predict the behavior of, for example, double mutants in the competitive FG phase assay from the behavior of the single point mutants. Deviations from ΔΔG linearity would point to coupling effects.

To explore this, we first combined positive mutations and indeed observed near-perfect additive effects (Fig. 4B, 4D). The A6C+K8L variant showed a ∼12-fold higher Xpo1 affinity – close to the product of the single mutation effects (2.6×5.3=14). N1W+K8L showed a 14.5-fold better binding, again close to the prediction (3.2×5.3=17). Thus, additivity applies here, and the PKI NES was brought to the level of a superNES by combining just two positive spacer mutations.

The data also show that the positive K8L mutation can compensate for the detrimental effect of the E4R substitution (Fig. 3B, 4B, 4D), yielding a tighter binder than the original PKI wild-type NES. In fact, the resulting double mutant bound Xpo1 2-fold better than predicted by a linear ΔΔG model, i.e. the K8L is even more beneficial in an E4R context than in a wild type one. As detailed in the Discussion, this mutational coupling is plausibly explained by the PKI NES·Xpo1 structure.

Similarly, we expected that combining negative spacer mutations would allow us to bring down the affinity for Xpo1 by three orders of magnitude, making the interaction physiologically irrelevant. To test this, we used a similar transfection-based setup as above but omitted the NLS from the reporters to test the negative spacer mutations pairs *in vivo*. The localization of the reporter then depends only on the balance between passive nuclear influx and export activity of the tested NES variant (Fig. 3D). Here, the wild-type PKI NES conferred full nuclear exclusion of the reporter, whereas an NES-free fusion remained evenly distributed throughout the cells.

The single A6K mutation clearly impaired the nuclear exclusion of the reporter, but still left it somewhat enriched in the cytoplasm. The E4R A6K, A6K A10T, and A6K N15E double mutations, however, completely inactivated the otherwise strong PKI NES, with the corresponding EGFP fusions showing an even nucleocytoplasmic distribution similar to that of the NES-free construct. Note that these loss-of-function mutants still perfectly conform to the consensus of a PKI-type NES with preferred residues at their Φ^1^–Φ^4^ positions.

### Engineering ‘nonNESs’

Having confirmed that the PKI NES point mutations can be combined with predictable effects, we wanted to apply this principle to the preference data for designing peptides with pre-defined properties in terms of Xpo1 binding from scratch. First, we wanted to generate sequences that look like PKI-type NESs but lack export activity. We designed three such sequences with leucines at Φ^1^–Φ^4^ (nonNESs1–3, Fig. 5A) as well as versions with a leucine also at Φ^0^ (nonNESs4–6). These sequences follow the consensus for a PKI-type NES, with the nonNESs4–6 even matching the 5Φ superNES pattern (Güttler *et al*., 2010). However, in each case, we included disfavored spacer residues and tested how these sequences affect the localization of EGFP fused to them (Fig. 5B). Notably, nonNESs1–4 reporters showed localization similar to that of EGFP without an NES indicating a complete lack of export activity. nonNESs5–6 may have retained some residual export activity but this was insufficient for nuclear exclusion of the reporter. Thus, 4 or even 5 optimal leucines at Φ positions are not sufficient for NES activity. Instead, they require a context of appropriate flanking and inter-Φ spacer residues.

**Figure 5.**
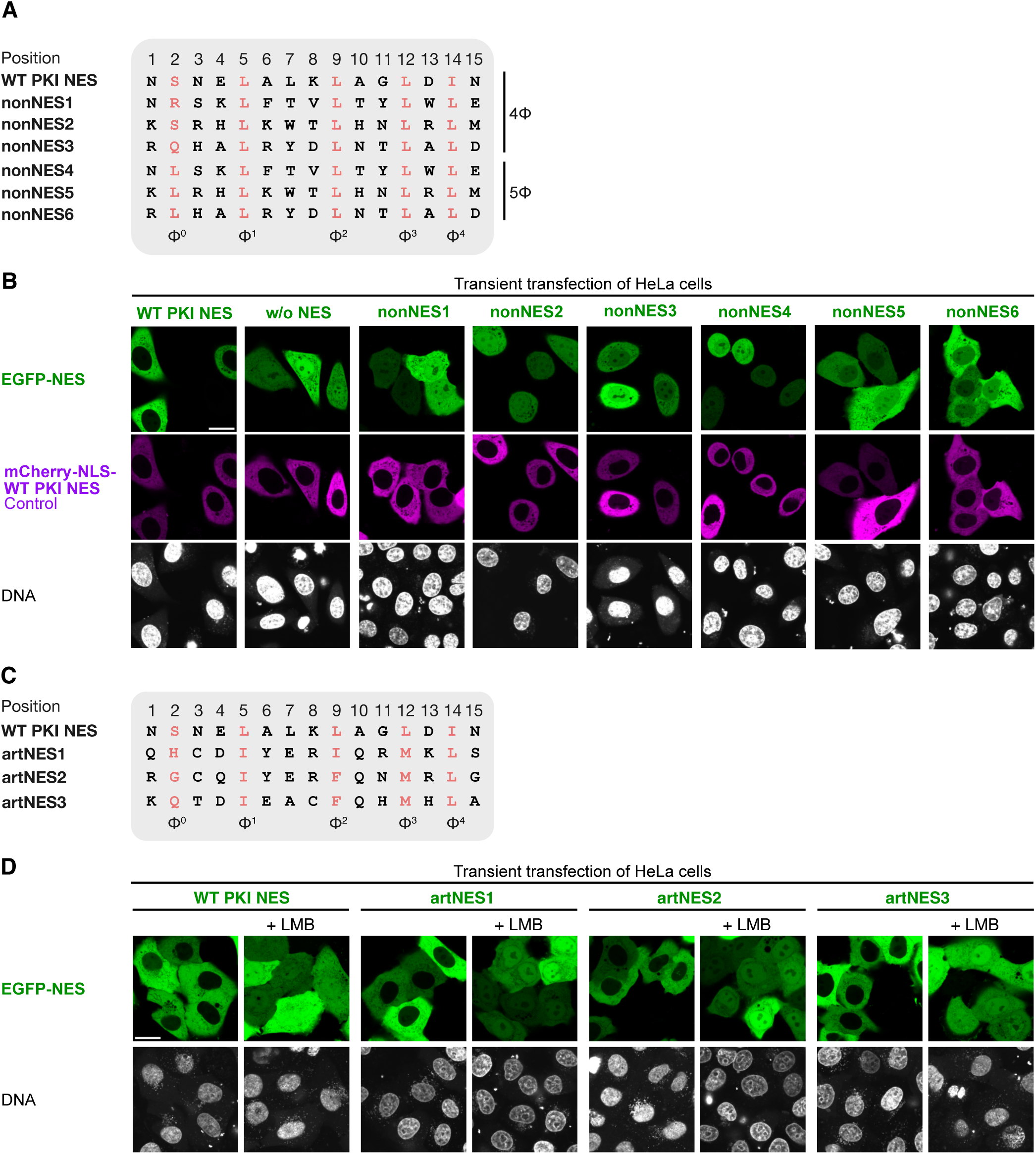
Engineered NES-like peptides. **(A)** Alignment of the PKI NES with engineered nonNESs conaining 4 or 5 leucines at the Φ positions but suboptimal spacers and flanks. **(B)** Localization of EGFP-nonNES fusions in cells, analyzed as in Fig. 3D. Despite matching the established NES consensus, none conferred nuclear exclusion. **(C)** Alignment of the PKI NES with artNESs — artificial NESs designed to mismatch the PKI sequence but to match its export activity. **(D)** Localization of EGFP-artNES fusions, as in (B), without and with LMB treatment (9 nM, 60 min). LMB sensitivity identifies the export as Xpo1-dependent.

### *De novo* NES design

Additionally, we set out to create artificial NES sequences (artNESs) with zero identity to the prototypic PKI NES but with similar export activity. To design such NES peptides, we selected residues for each position according to the amino acid preferences (Fig. 1B) such that they were different from those in the corresponding positions of the PKI NES but matched their preference values as closely as possible (Fig. 5C). The designed sequences notably deviated from the common scheme of a leucine-rich NES, since all the positions but Φ^4^ were occupied by residues other than leucine. We tested three such artNESs as EGFP fusions in the transfection assay and observed perfect LMB-sensitive nuclear exclusion for all of them (Fig. 5D). Thus, the phage display-derived preference dataset can guide the *de novo* design of functional Xpo1-dependent NESs.

### Engineering ‘hiNESs’

The next step was to design ‘hiNESs’ – peptides with the highest possible Xpo1 affinity – by filling each NES sequence position with the most preferred residue. Implementing this, however, yielded a highly aggregation-prone peptide (aggNES). Although it still binds Xpo1, it does so with a lower affinity than expected (see below), one possible reason being that only a small non-aggregated fraction remains available for exportin binding. The corresponding EGFP-aggNES fusion showed the expected nuclear exclusion, but also a prominent Golgi mislocalization (Fig. 6B) – an off-target interaction. Thus, aggNES retained Xpo1 binding but lost its targeting specificity.

**Figure 6.**
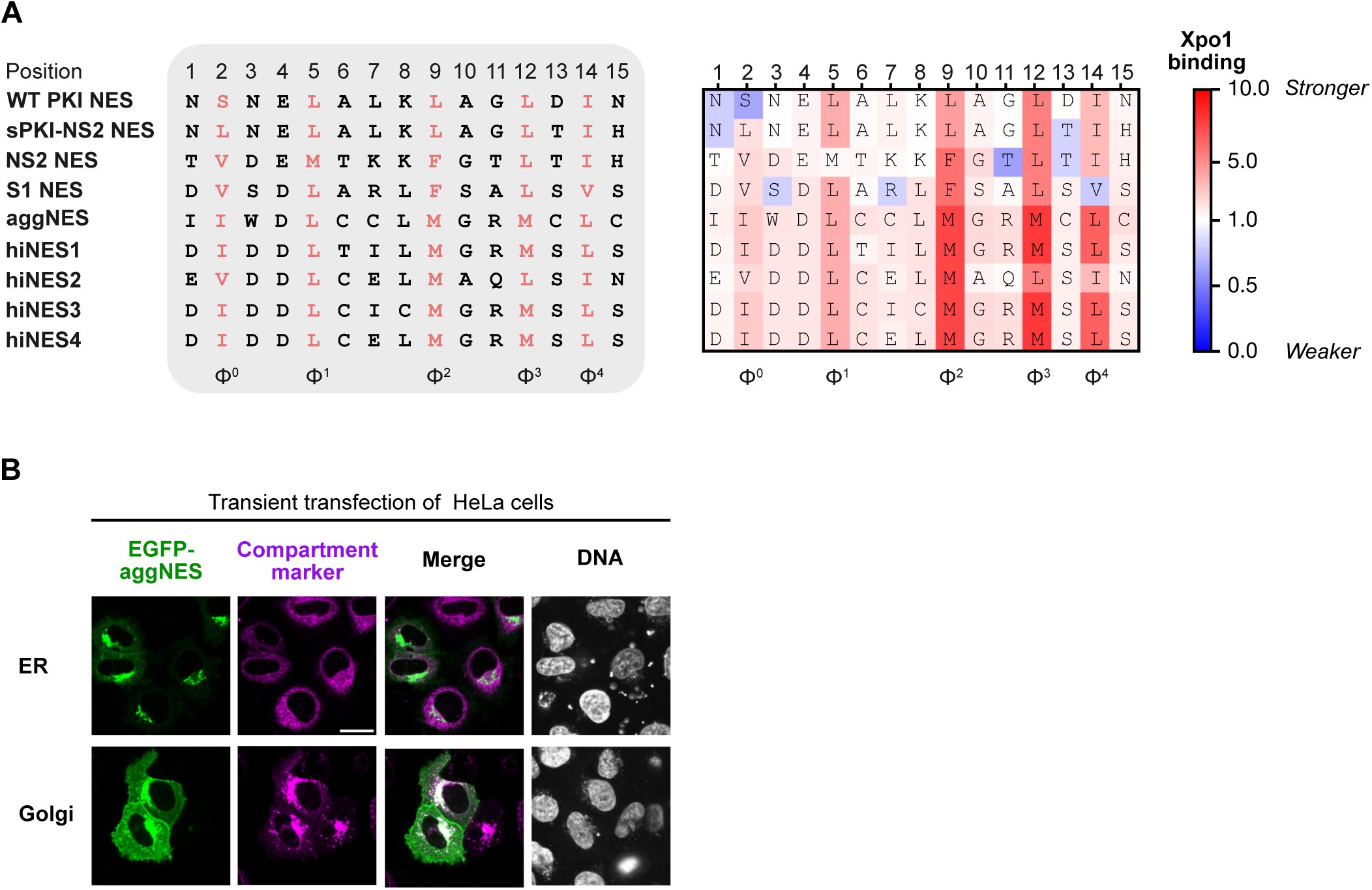
hiNES design. **(A)** Alignment of the PKI NES with previously described superNESs and with the hiNESs, together with a heatmap of the corresponding human Xpo1 preference values (Fig. 1B). **(B)** Cellular localization of EGFP-fused aggNES, which represents a failed hiNES design with off-target localitzations. The fusion is excluded from the nucleus, indicating export activity, but also accumulates in bright structures that co-localize with a Golgi marker (Golgi-7; Addgene #55052) but not with an ER marker (mCherry with an N-terminal signal sequence and a C-terminal ER retrieval signal; Munro and Pelham, 1987). Some plasma membrane staining is also evident. Scale bar, 20 µm.

What is the issue here? Hydrophobic residues are preferred not only at the Φ positions, but also at several spacer positions – for example, tryptophan flanking Φ^0^, isoleucine and leucine between Φ^1^ and Φ^2^, and cysteine throughout (Fig. 1B-C). Filling every position with its most preferred residue thus turned seven of the ten spacer and flanking residues hydrophobic (four Cys, one Trp, one Ile, and one Leu), yielding an overly hydrophobic, poorly soluble peptide with off-target interactions.

For the following designs, we therefore treated water solubility as a key constraint, balancing the hydrophobic-hydrophilic ratio and favoring hydrophilic residues wherever the preference difference was small (e.g., acidic residues over tryptophan flanking Φ^0^). hiNES1 was built free of cysteines for oxidation resistance and maleimide-labeling compatibility; hiNES2 and hiNES3 carried one and two cysteines, respectively; and hiNES4 was designed as a hiNES1/hiNES2 chimera (Fig. 6A).

We then asked how the Xpo1 affinities of the newly engineered hiNESs compared with those of previously reported ‘superNESs’, namely the synthetic S1 NES (Engelsma *et al*., 2004) and the naturally occurring viral NS2 NES (Engelsma *et al*., 2008) — both including the initially overlooked additional Φ^0^ valine that contributes to their high affinity (Güttler *et al*., 2010) and, finally, the sPKI-NS2 NES chimera (Fu *et al*., 2018) A comparison with the preference heatmap (Fig. 6A) revealed that all of these superNESs have two to three positions occupied by disfavored residues and several more by suboptimal ones. In contrast, the hiNESs have optimized sequences without any disfavored residues. Moreover, all of the hiNESs have more optimal sets of the Φ residues.

To measure the affinities of the abovementioned superNESs and hiNESs to Xpo1, we labeled NES-ZZ-fusions via an ectopic cysteine several residues downstream of the NES with fluorescein maleimide (Güttler *et al*., 2010) and measured fluorescence anisotropy after incubation with varying Xpo1 concentrations in the presence of a saturating RanGTP concentration (Fig. 6B). The K_D_ of 2.6 nM measured for the NS2 NES agrees well with the previously reported value (Fu *et al*., 2018). For the sPKI-NS2 NES chimera, we observed a K_D_ of 7.1 nM, placing this chimera between the NS2 NES and the PKI NES Φ^0^L. The estimated K_D_s for S1 NES and hiNES1 were at or below 1 nM, which is outside the reliable range for this method (considering that 10 nM concentration of labeled peptides was used for a high signal-to-noise fluorescence signal).

To be able to discern the affinities of the high-affinity superNESs, we also performed fluorescence anisotropy measurements in the absence of RanGTP (Fig. 6C). This reduces the affinity of the NES•Xpo1 interaction by three orders of magnitude (Güttler *et al*., 2010). Indeed, we could barely observe any change in anisotropy for the NS2 NES and sPKI-NS2 NES even at 1 µM Xpo1. For the S1 NES, binding became notable at ≥ 250 nM Xpo1, but the data were not sufficient to reliably extract a K_D_. On the other hand, a K_D_ of 110 nM was estimated for the interaction of hiNES1 with Xpo1 in the absence of RanGTP. This is comparable to the affinities of many physiological NESs in the presence of RanGTP (Fu *et al*., 2018) and similar to the cellular Xpo1 concentration of 100–200 nM (Wühr *et al*., 2014; Kirli *et al*., 2015), predicting a substantial engagement with the exportin even under cytoplasmic conditions.

To extend measurements of NES affinities for Xpo1 (in the presence of RanGTP) to the sub-nanomolar range, we applied a competitive setup to our anisotropy measurements, first using the fluorescently labeled NS2 NES fusion as a constant reference and varying the concentrations of a given NES-MBP competitor as the analyte (Fig. 7A). This yielded essentially the same ranking as the direct assay but with approximately 2-fold lower K_D_s (perhaps because of a bias introduced by the particular fluorophore-peptide linkage used here). The values were 4 nM for sPKI-NS2 NES, 1.1 nM for the NS2 NES, and a 5-fold better affinity (K_D_ of 0.2 nM) for the S1 NES.

**Figure 7.**
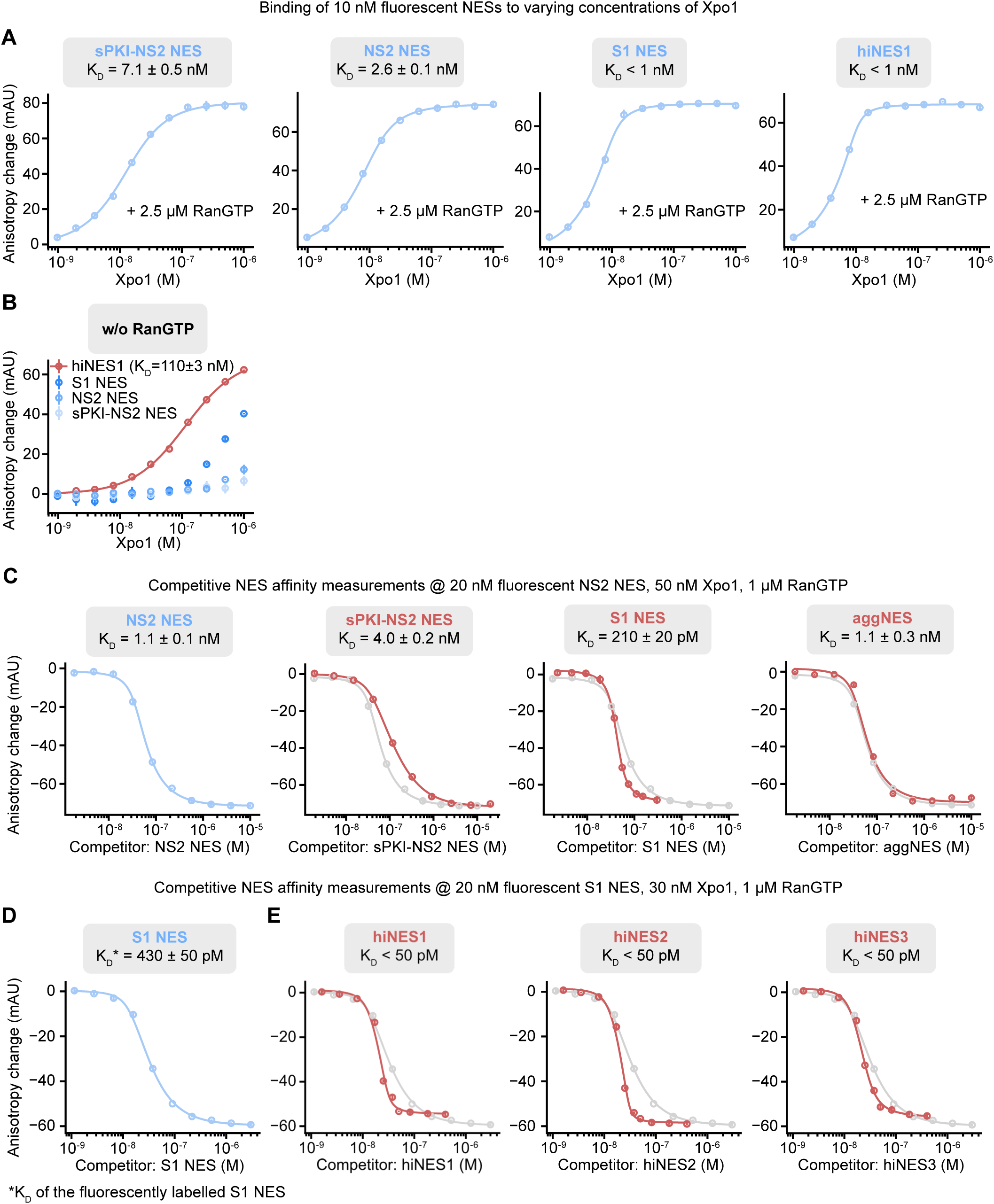
Affinity measurements for superNESs and hiNESs by fluorescence anisotropy. **(A)** Anisotropy of 10 nM fluorescently labeled peptides titrated with hXpo1 in the presence of excess hRanGTP(5–180, Q69L). Bound peptides show higher anisotropy. K_D_s were obtained by fitting a direct-binding model. Error bars, standard error of technical triplicates. **(B)** As in (**A**) but without hRanGTP. Fitting was reliable only for hiNES1 that showed the strongest binding. **(C)** Competition experiments: 20 nM fluorescently labeled NS2 NES-ZZ with 50 nM hXpo1 and excess hRanGTP(5–180, Q69L), titrated with MBP-fused competitor peptides. K_D_s of the competitors were obtained by fitting a competitive-binding model. The NS2 NES-MBP data are shown separately in blue and, for reference, in gray on the other graphs. Error bars, standard error of technical triplicates. **(D)** As in (**C**), but with 20 nM labeled S1 NES-ZZ and 30 nM hXpo1, titrated with S1 NES-MBP. The K_D_ of the labeled S1 NES was obtained with the competitive-binding model, using the K_D_ of S1 NES-MBP from (**C**); the asterisk in the panel marks this value. **(E)** As in (**D**), but with hiNES-MBP competitors. The S1 NES binding curve from (D) is shown in gray for reference.

To attempt to measure the Xpo1 affinities of the hiNESs (in the presence of RanGTP), we switched to a fluorescently labeled S1 NES as a reference because it already showed a tighter Xpo1 interaction than the NS2 NES. This revealed that the apparent K_D_s are under 50 pM, with hiNES3 displaying somewhat weaker affinity than hiNES1 and hiNES2 (Fig. 7C). A more precise estimation was not possible even in this competitive setup with the high-affinity S1 NES reference: here, too, the assay reached its limit. Nevertheless, all the hiNESs clearly bind more tightly than the S1 NES.

The competitive assay also indicated that aggNES binds Xpo1 with a K_D_ of only ∼ 1 nM (Fig. 7A, C), which is at least 20-fold weaker than hiNES2. This assay provides only a lower bound on the hiNES2 affinity. Indeed, a more appropriate assay (Fig. 8 and Table 1) suggests an even larger factor of ∼50-fold. Given that the aggNES sequence adheres most closely to the preference dataset, this is a striking underperformance. One explanation is that aggregation competes with exportin binding (as alluded to above) — though aggregation might not persist at the low peptide concentrations used here. In addition, an incompatibility (negative positional coupling) between the singly preferred residues likely reduces affinity here. We detail this phenomenon below for hiNES1, hiNES2, hiNES3, and hiNES4, as these variants can be compared without the complication of aggregation effects.

**Figure 8.**
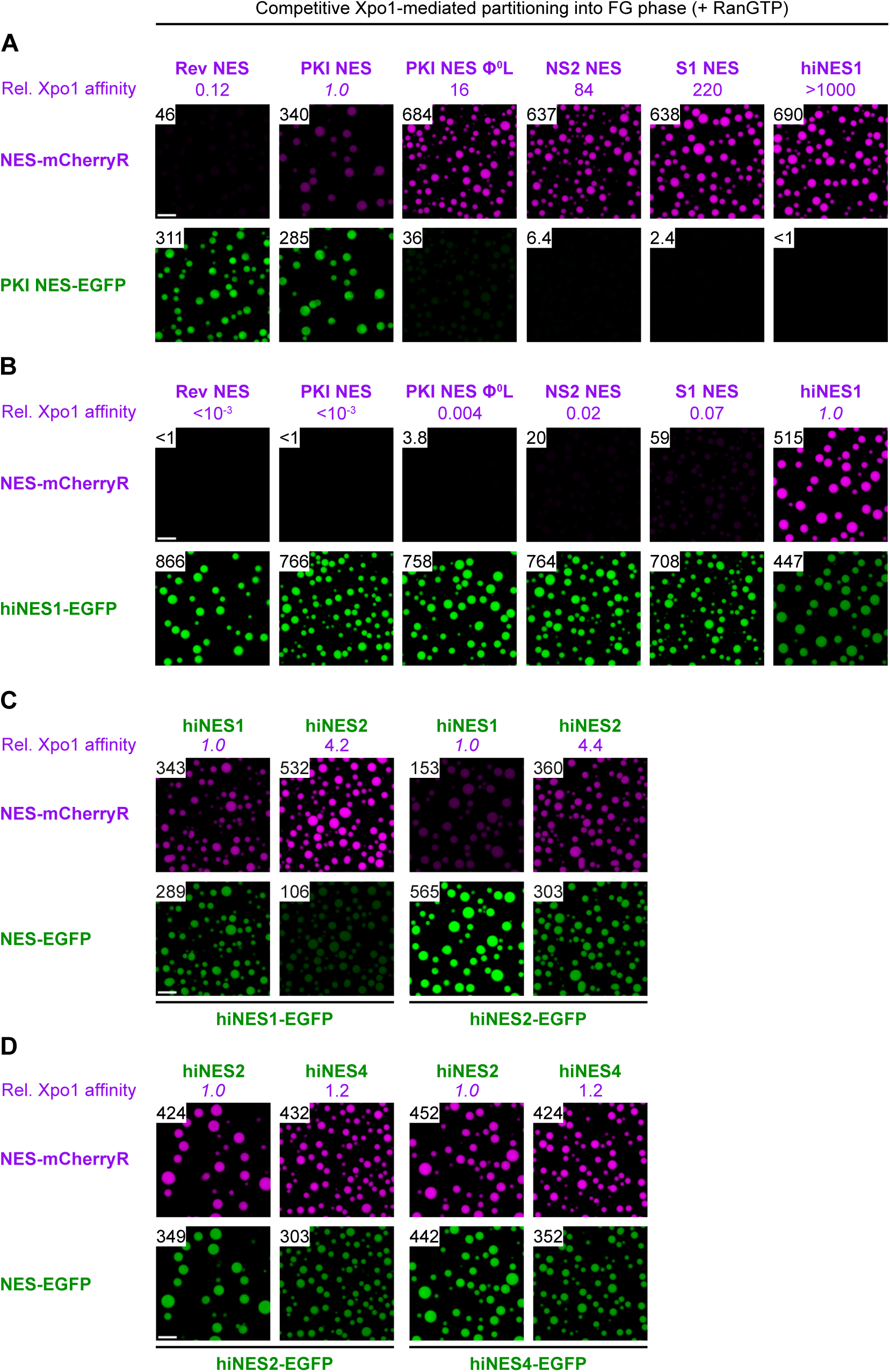
Extreme Xpo1 affinities of the hiNESs revealed by chained, wide-range FG phase competition. **(A)** A series of NESs (fused to mCherryR) competed with the WT PKI NES-EGFP standard for limiting Xpo1 in an FG partitioning assay (as in Fig. 3B). From Rev NES up to NS2 and S1, the competitors span a wide affinity range but can still be measured against the standard. hiNES1, however, competed so strongly that the PKI standard remained phase-excluded, so that a reliable ratio of partition coefficients could not be derived. **(B)** Similar experiment as in (**A**), the difference being hiNES1-EGFP as the constant competitor, so that Xpo1 affinities are given relative to hiNES1. Comparing (**A**) and (**B**) then allows the affinity of hiNES1 to be estimated relative to PKI wild type, by ‘chaining’ through intermediate binders (PKI NES Φ⁰L, NS2, S1): hiNES1 binds 3700-fold more strongly than the WT PKI NES. **(C)** hiNES1 and hiNES2 competed pairwise, each measured in both directions (as EGFP and as mCherryR fusion); the two directions gave the same ratio. **(D)** As in (**C**), for hiNES2 and hiNES4. Together with (**A–C**), this chains the hiNES4 affinity onto the PKI scale, giving a ∼20,000-fold higher affinity than the WT PKI NES. Combined with the 16 nM affinity measured for the PKI NES Φ⁰L NES, this places hiNES4 at a ∼10–15 pM K_D_ for the Xpo1·RanGTP subcomplex. Scale bar, 10 µm.

The extreme affinities of the hiNESs lie already well beyond the reliable range of the competitive anisotropy assay (even using the S1 NES reference); the derived K_D_ values are therefore only approximations. We therefore turned back to our competitive FG phase partitioning assay, upgrading it to extend the measurement range.

The need for such an upgrade is illustrated by the observation that the hiNESs outcompete our wild-type PKI NES standard to the point of phase exclusion (Fig. 8A). This precludes a proper ratiometric quantification and renders a comparison of different high-affinity binders through this PKI standard uninformative.

Indeed, comparing a wide range of NES affinities requires more than one reference point. We therefore developed an entire series of calibrated NES-mCherryR standards, spanning five orders of magnitude in Xpo1 affinity. The panel includes the weaker physiological Rev NES, the stronger physiological PKI NES, the high-affinity PKI Φ^0^L, NS2, and S1 NESs, as well as the newly engineered hiNES1, hiNES2, and hiNES4 variants (Fig. 8A-D, and Table 1). To assess robustness, we included two-way calibrations, comparing X/Y NES pairs interchangeably as EGFP-X/mCherryR-Y and EGFP-Y/mCherryR-X pairs, which yielded very consistent ratiometric affinities.

On the low affinity end, the data indicate that the Rev NES binds Xpo1 8-fold weaker than a wild-type PKI. On the high affinity end, we observed the following increases in Xpo1 affinity over the PKI standard: PKI Φ^0^L (16-fold), NS2 (84-fold), S1 (220-fold), hiNES1 (3700-fold), hiNES2 (16000-fold), and hiNES4 (19000-fold).

It was initially surprising that hiNES2 binds human Xpo1 more strongly than hiNES1, given that hiNES2 is less optimal at Φ^0^ and spacer position 7 (Figs. 1B, 6; Table 1). In fact, however, this points to a neighboring effect, or more specifically to negative coupling between the most preferred residues within the α-helical Φ^1^–Φ^2^ spacer: the β-branched residues in the Thr-Ile (-Leu) motif of hiNES1might impose steric constraints on the α-helical conformation required for Xpo1 binding. The same applies to hiNES3, where this spacer harbors a Cys-Ile-Cys motif, containing a β-branched Ile and two adjacent Cys residues (with bulky thiol moieties), which have a similarly low α-helical propensity as threonine (Pace and Scholtz, 1998). In contrast, the Cys-Glu-Leu spacer in hiNES2 appeared more optimal in this respect, given that Glu and Leu are less bulky near the polypeptide backbone. With this in mind, we transplanted the hiNES2 spacer onto the hiNES1/3 background to generate hiNES4 (Fig. 6A). Indeed, the competitive FG phase-partitioning assay revealed that hiNES4 binds Xpo1 5-fold tighter than hiNES1, 1.2-fold tighter than hiNES2, and ∼19,000-fold tighter than the wild-type PKI NES (Fig. 8C–D). This qualifies hiNES4 as the highest-affinity NES described to date (Table 1).

### Structural characterization of the hiNES2 interaction with Xpo1

To gain insight into the structural basis for the high affinity of the hiNESs, we crystallized hiNES2 in complex with Xpo1 and RanGTP and solved the structure at 3.0 Å resolution (PDB: 8QYZ; see Methods and Appendix Table S4 for details and statistics). We chose Xpo1 and Ran from *S. cerevisiae* because these were co-crystallized with the PKI NES before (Koyama *et al*., 2014), avoiding the need for a snurportin 1 chimera (Güttler *et al*., 2010) that would otherwise bias the orientation of the bound NES. Furthermore, hiNES2 was designed according to the yeast Xpo1 preferences, including the Leu-over-Met preference at Φ^3^.

As predicted, hiNES2 docks into the peptide-binding cleft of Xpo1 in the same orientation as the PKI NES (Fig. 9A). Furthermore, all predicted Φ residues of hiNES2 occupy the corresponding hydrophobic pockets of the NES-binding site (Fig. 9B, C). Given that hiNES2 contains other hydrophobic residues, this is not a trivial outcome. Altogether, this indicates that hiNES2 conforms to the PKI NES class not only in sequence, but also in binding mode.

**Figure 9.**
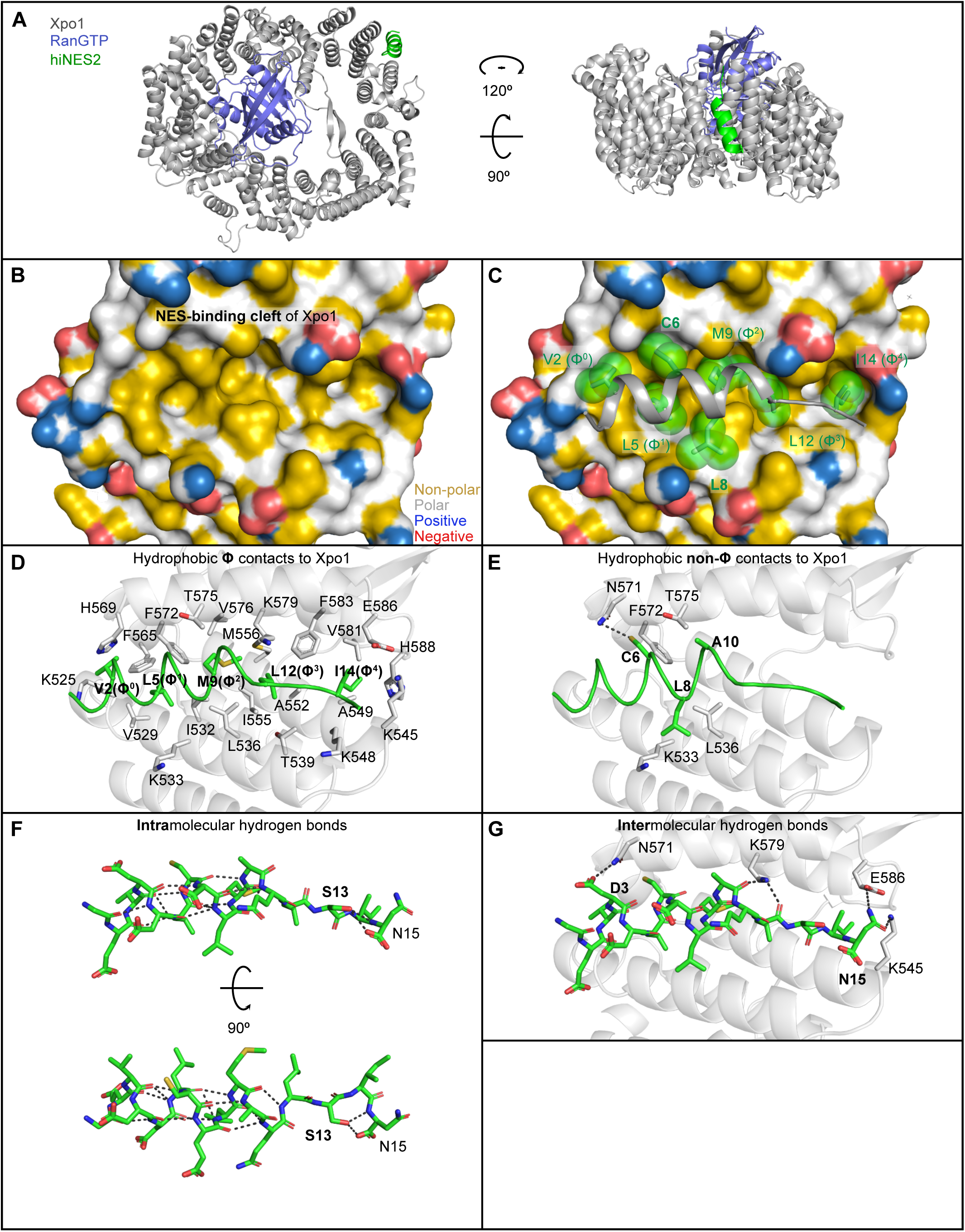
Crystal structure of the hiNES2·yXpo1·RanGTP complex. (**A**) Overall structure, shown as a cartoon in two orientations. yXpo1, gray; RanGTP, blue; hiNES2, green. (**B**, **C**) Hydrophobicity of the hiNES2-binding pocket of yXpo1, with hiNES2 omitted (**B**) or shown as a cartoon (**C**). hiNES2 residues that bury more than 50 Å² of their own surface are drawn as sticks and semi-transparent spheres (side chains and Cα); hiNES2 is green. Φ residues are labeled; hydrophobically interacting spacer residues are in bold. yXpo1 is shown as a surface, colored by the YRB hydrophobicity scheme (Hagemans et al., 2015). (**D**, **E**) Hydrophobic contacts of the Φ-residue side chains (**D**) and non-Φ-residue side chains of hiNES2 with yXpo1. yXpo1, gray cartoon; hiNES2, green ribbon, with the relevant side chains (and Cα) as sticks. yXpo1 side chains within 5.0 Å of these hiNES2 side chains (and their Cα) are shown as sticks and labeled. In (E), L8 packs against K533 and L536, A10 against T575, and a hydrogen bond between yXpo1 N571 and hiNES2 C6 is drawn as a black dashed line. **(E)** Intramolecular hydrogen bonds within hiNES2 (Xpo1 omitted), in two orientations. hiNES2 is shown as green sticks; hydrogen bonds as black dashed lines. Note the bonds from the S13 side chain to the backbone amide of N15. **(F)** Intermolecular hydrogen bonds between hiNES2 and yXpo1. yXpo1, gray cartoon; hiNES2, green sticks; hydrogen bonds, black dashed lines. Residues whose side chains form the bonds are labeled — in bold for hiNES2; for yXpo1, their side chains and Cα are shown as sticks. Some of these bonds vary between the four complexes in the asymmetric unit.

As expected, hiNES2 primarily relies on hydrophobic interactions for binding to Xpo1 (Fig. 9B-E). Its Φ residues account for 596 Å^2^ of the 892 Å^2^ buried at the interface (values refer to the hiNES2 side). Φ^2^ M9 appears to be a particularly strong contributor. It packs against Xpo1: M556, F572, and V576 and buries 119 Å². Its flexible sidechain adopts a shape more complementary to the pocket than a leucine’s, allowing more extensive van der Waals contacts. This rationalizes the preference for methionine at Φ² in both human and yeast Xpo1 (Fig. 1B, C).

Outside the Φ residues, the sidechain of spacer L8 packs against K533 and L536 of Xpo1. It buries only moderately less surface than in Φ-positions leucines (73 vs 112-123 Å^2^), so it likely contributes to the overall stability of the interaction, explaining why leucine is strongly favored at position 8 (Fig 1B, C) and why K8L has such a strong Xpo1 affinity-enhancing (Fig. 4B). The A10 sidechain packs against the methyl group of T575 of Xpo1 as in the PKI NES Φ^0^L structures (Güttler *et al*., 2010; Koyama *et al*., 2014).

Furthermore, the hiNES2 C6 sidechain is accommodated by a hydrophobic cavity at the top of the NES-binding cleft (Fig. 9C, E), burying 64 Å^2^ and packing against Xpo1: F565 and N571. It fills this cavity more completely and makes more extensive van der Waals contacts than the corresponding alanine at the same position of the PKI NES. Additionally, the thiol group of C6 can form intermolecular hydrogen bonds. Although hydrogens are not visible at our resolution, and the four complexes in the asymmetric unit show some sidechain flickering in this region, the amide group of N571 is close enough to allow either N–H···S or O···H–S contacts.

The dual nature of cysteine, capable of forming both hydrophobic and polar contacts, explains why it is favored at this position and well tolerated elsewhere in the NES spacers. The complex is further stabilized by a few other intermolecular hydrogen bonds between sidechains (Fig. 9G), such as between hiNES2: D3 and Xpo1: N571, and between hiNES2: N15 and Xpo1: E586 or K545 (again with some sidechain flickering between the complexes).

Finally, the negatively charged residues flanking Φ^0^ are electrostatically attracted to the positively charged surface near the NES-binding pocket of Xpo1(Appendix Fig. S1D). This charge complementarity aligns not only with our preference data (Fig. 1) but also with previous findings that a negative charge at the N-terminal part of an NES enhances its affinity for Xpo1 (Güttler *et al*., 2010).

hiNES2 adopts a characteristic secondary structure comprising an α-helix transitioning into an extended conformation (Fig. 9A, C), with an uninterrupted intra-helical hydrogen-bonding network. This resembles the NS2 NES (Fu *et al*., 2018) but contrasts with the PKI NES Φ^0^L, which lacks some of these bonds (Güttler *et al*., 2010; Koyama *et al*., 2014). The sidechain of S13 forms intramolecular hydrogen-bonds to the backbone amide of N15 (Fig. 9F) and thereby stabilizes the bound conformation. This explains why serine is preferred at position 13 (Fig. 1B, C).

The favorable effect of these additional intra-NES hydrogen bonds on the hiNES2–exportin interaction can be described in three equivalent ways: as a stabilization of the bound conformation against dissociation, as a reduced entropic penalty for capturing an otherwise flexible peptide, or as a shift of the free-peptide equilibrium toward the binding-competent state (conformational selection).

### Behavior of hiNESs in nuclear transport

Finally, we examined the *in vivo* effects of the exceptional Xpo1 affinity displayed by hiNESs. We transiently expressed EGFP-fused hiNESs and other superNESs in HeLa cells alongside an export control (mCherry-SV40 NLS-PKI NES). We used the PKI NES reporter as a baseline for a strong, physiological NES; it was excluded from nuclei and did not impair export of the co-expressed control (Fig. 10A). After LMB treatment, the PKI NES reporter redistributed evenly throughout the cells, whereas the NLS-containing control accumulated inside nuclei.

**Figure 10.**
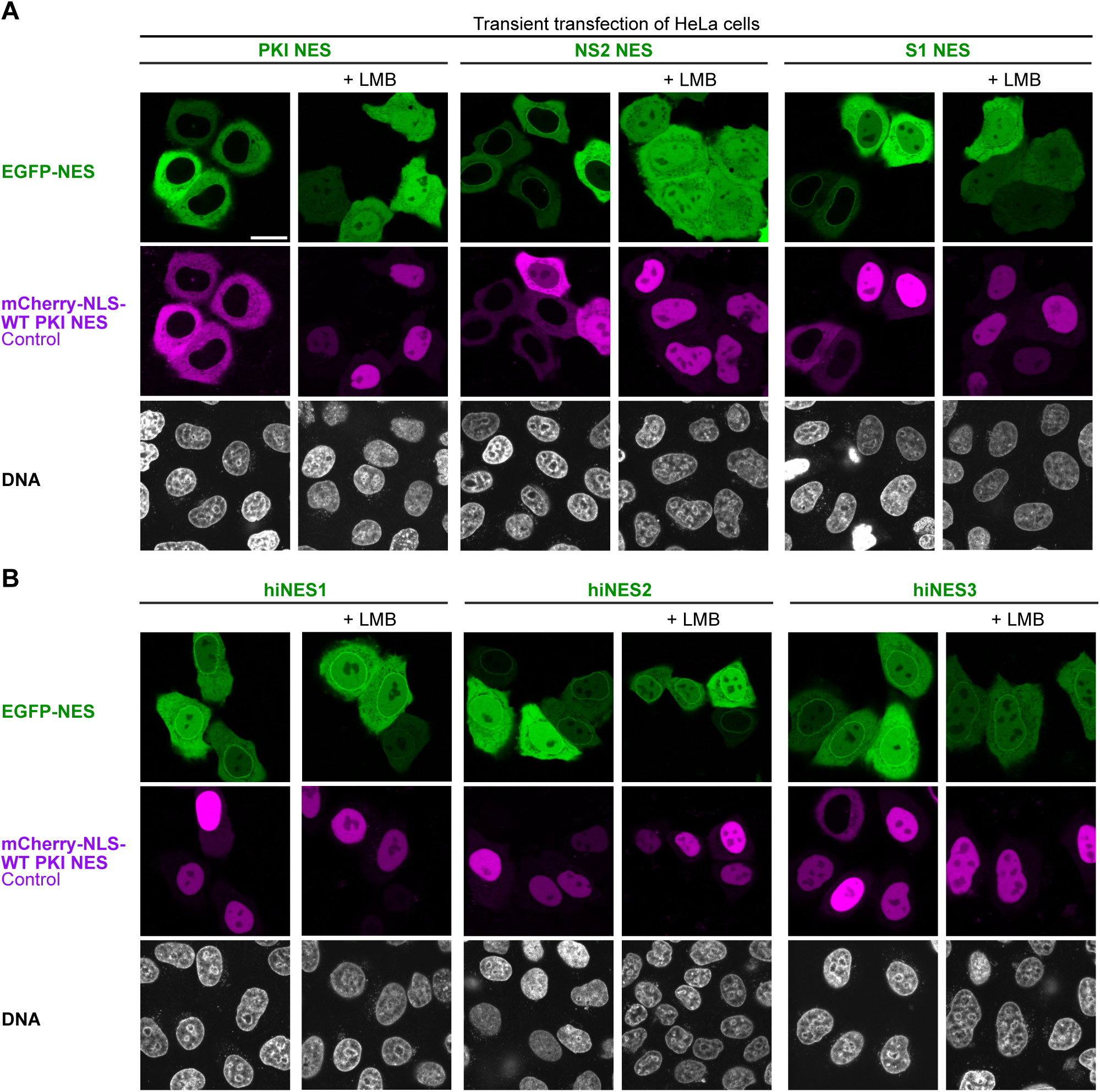
hiNESs jam Xpo1 irreversibly and fail to confer export. **(A)** The indicated NES-EGFP fusions were transiently expressed in HeLa cells. The PKI NES shows LMB-sensitive nuclear exclusion and allows export of the mCherry control. NS2 and S1 also show normal export at low expression. At higher expression, however, NS2 mislocalizes the mCherry-NLS-WT PKI NES control reporter, whereas S1 disrupts both the control and its own export. Both show NPC staining that disappears upon LMB treatment (9 nM, 90 min), indicating still-dynamic (and thus competable) Xpo1 interactions. **(B)** The hiNESs, tested as in (**A**). None are excluded from the nucleus, and all block export of the control reporter. All give crisp NPC staining that resists LMB treatment. This indicates an irreversible hiNES–Xpo1 interaction, given that LMB blocks the loading of NESs onto Xpo1 but cannot displace the already-formed hiNES·Xpo1 complexes. Scale bar, 20 µm.

NS2 NES-EGFP exhibited perfect, LMB-sensitive nuclear exclusion. In addition, a (weak) nuclear rim signal was evident, which likely originates from NPC-bound transport intermediates and points to slow release from Xpo1 (Engelsma *et al*., 2004, 2008). In cells with higher NS2 NES-EGFP expression, nuclear exclusion of the co-expressed export control was lost, indicating a competitive block of Xpo1. The S1 NES fusion behaved similarly, though its own nuclear exclusion was complete only in weakly expressing cells; at higher expression levels, nuclear exclusion was abolished, indicating self-competition and saturation of the export pathway.

Cells expressing the hiNES1–3 reporters displayed several unique features (Fig. 10B). First, these fusions distributed evenly between the nucleus and cytoplasm, even at low expression levels. The only site of enrichment was the nuclear rim (NPCs). This indicates that hiNESs do not confer unidirectional export but fail to dissociate from the exportin in the cytoplasm, causing Xpo1 to shuttle them bidirectionally. Second, the PKI NES-based export control equilibrated into nuclei in all cells expressing a hiNES reporter, demonstrating that hiNES expression blocks Xpo1 as effectively as LMB treatment. Third, the localization of the hiNES reporters at the nuclear rim was remarkably resistant to LMB, persisting even after a 90-minute treatment. This documents an essentially irreversible interaction with Xpo1 under *in vivo* conditions and is consistent with structural data showing that a bound NES masks C528, preventing the Michael addition of LMB.

Such persistence implies off-rates on the order of at least hours. This would be consistent with the 15–70 pM affinities of the hiNESs for the Xpo1·RanGTP species (assuming on-rates on the order of 10^4^–10^5^ M^-1^s^-1^), but hard to reconcile with their 1000-fold weaker affinities for the exportin alone (110 nM for hiNES1). The most likely explanation is that Xpo1 is not switched to its Ran-free, low-affinity state — the state from which an NES would normally be released.

## Discussion

Binding of short linear motifs by proteins underlies many biological processes, including gene expression (Zaborowska *et al*., 2016), protein sorting (Görlich and Kutay, 1999; Teufel *et al*., 2022), and immunity (Pishesha *et al*., 2022). Understanding the rules behind such interactions is therefore widely desirable. Yet even for short peptides the sequence space is vast, and predicting which sequences bind a given protein, and how well, remains challenging.

We present here a general strategy for this problem. A phage-display library samples the sequence space, and a selection retrieves binders; because each peptide variant is physically linked to its coding sequence as a barcode, deep sequencing can count and normalize binding events, upgrading the workflow to a quantitative, high-throughput affinity ranking. Applied to PKI-class NESs, it yielded a comprehensive position-specific preference map for the Xpo1 interaction — reproducing the known Φ preferences and making them quantitative, and revealing that the inter-Φ spacer positions, typically written as “x” in consensus definitions, are anything but neutral: they strongly shape Xpo1 affinity.

Two engineered sets illustrate this in opposite directions. The nonNESs (Fig. 5A–B) satisfy the classical leucine-rich rule (with leucines at the Φ positions) yet fail to confer export — correct Φ anchors in a poor spacer context do not make an NES. Conversely, the hiNESs reached picomolar affinity only by optimizing the Φ and spacer positions together. The spacers thus carry much of an NES’s functional information, not by fixing it to one sequence but by grading the many anchor-correct sequences from non-functional to optimal.

Building on earlier work on FG hydrogels (Frey and Görlich, 2007; Schmidt and Görlich, 2015; Ng *et al*., 2021), we developed a ratiometric assay to quantify NES–Xpo1 interactions across an exceptionally wide affinity range (micromolar to low picomolar), in a setting that recapitulates the chemical environment of nuclear pore passage. It measures the competition of differently colored NESs for Xpo1 binding and uses the partitioning of the export complexes into an FG phase as a readout. This approach extends naturally to other NTR–cargo interactions — whether signal-, conformation-, or adaptor-based — to pinpoint cargo features relevant for recognition, and more broadly to client interactions in other phase-separated compartments (Musacchio, 2022) whose roles in physiology and pathology are increasingly evident (Alberti and Hyman, 2021).

The phage-display and deep-sequencing pipeline calls for broader application, for example to map the peptide-binding preferences of other proteins. It could also identify ideal binders against a given target from a combinatorial library, or affinity-mature sub-optimal variants from a mutagenized library. This overcomes a fundamental limitation of traditional display techniques, namely that physical libraries sample only a tiny fraction of the relevant sequence space. Crucially, the optimal binder need not be present in the library at all: it is assembled computationally by taking the most-preferred residue at each variable position and correcting for positional couplings. This holds great promise for developing antibodies, nanobodies, or alternative scaffolds such as affibodies (Nord *et al*., 1997) or DARPins (Forrer *et al*., 2003). As a proof of concept, our best hiNES binds Xpo1 ∼1,000-fold more tightly (K_D_∼13 pM) than the best binder present in the starting library (K_D_∼16 nM). Finding it required only a positional scan of ∼300 variants (15 positions × 20 choices), each quantified by deep sequencing. A direct selection of the optimum would instead have demanded a library of 20^15^.

Ultimately, one would like to predict NES activity from sequence. Several predictors exist (la Cour *et al*., 2004; Fu *et al*., 2011; Kosugi *et al*., 2014; Prieto *et al*., 2014; Xu *et al*., 2015; Engler *et al*., 2026), but they do not adequately consider the impact of the inter-Φ spacers — they attribute high scores, for example, to the nonNESs described here, which in fact fail to confer export. It is now tempting to use our preference datasets for more accurate predictions of Xpo1 affinity, and ultimately of NES activity. This would involve scanning candidate protein sequences in a sliding window of 15 residues and calculating a local score by multiplying the positional preference values, which we provide as a Scoring Matrix in the Appendix, combining the data from this study with the previously published data (Güttler *et al*., 2010) The multiplication gives a product that scales with the expected affinity. It corresponds to summing up ΔG increments. There are, however, still several limitations and caveats.

First, although the PKI-type accounts for the vast majority of NESs, other Xpo1-binding modes are not adequately described by the PKI preference list: the snurportin-type NES (with a Φ^2^-xxx-Φ^3^ instead of a Φ^2^-xx-Φ^3^ pattern), the Rev NES, which binds in a non-helical conformation, and the reverse NES. This list is possibly not even complete. Before comprehensive NES prediction becomes feasible, these gaps must be closed.

Second, positional preferences are not strictly independent: one position’s preference is influenced by its neighbors, and the multiplicative model is therefore only an approximation. This is most evident among the hiNESs (Table 1). hiNES3 attains the highest prediction score of all (1750), yet binds most weakly of the four based on the fluorescence anisotropy measurements, whereas hiNES2 has a lower score (112) and still binds an order of magnitude more tightly than hiNES1 (16 pM vs 70 pM). This inverted ranking can be attributed to the Φ^1^– Φ^2^ spacer, whose individually most-preferred residues (Cys, Thr, Ile) share a low α-helical propensity, so that their co-occurrence penalizes the very conformation the NES must adopt for binding (see above). Exchanging this spacer accordingly improved affinity 5-fold (hiNES4, K_D_ 13 pM, versus hiNES1, 70 pM).

The non-linear-additivity of the E4R and K8L mutations (Table 1, Figs. 3B, 4B, 4D) illustrates that couplings also arise between residues that are distant in sequence but adjacent in the bound conformation. Positions 4 and 8 face each other across one helical turn, flanked by Lys522 of human Xpo1 and (∼5 Å away) by Glu529. Lys8 of the wild-type NES is electrostatically neutral there — repelled by the former, compensated by the latter — so that K8L is beneficial simply because leucine adds a hydrophobic contact (K_D_ 260 → 48 nM, a 5-fold gain). On an E4R background this hydrophobic gain persists, but Arg4 introduces a further positive charge next to Lys522, so that Lys8 is now electrostatically unfavorable. K8L therefore also removes an unfavorable charge, and its effect increases correspondingly, to ∼9-fold (K_D_ 1.1 µM → 120 nM).

Electrostatic effects can also explain why the S1 NES binds far more tightly than its low score (4.8) would suggest. Its arginine in position 7 is rather disfavored in a PKI context (in which the preferences were measured) because of its vicinity to Lys8. S1, however, already carries the compensatory K8L exchange that removes that repulsive charge. The contribution of a residue thus depends on its neighbors — whether mediated by backbone conformation or electrostatics, within the peptide or with the receptor — which amounts to epistasis between positions and cannot be captured by a purely position-wise score. While most of such coupling effects can be plausibly explained by structural models, anticipating and enumerating all combinatorially possible couplings is out of reach.

Third, the candidate peptide must be accessible: even a high-scoring stretch binds Xpo1 only if solvent-exposed and not buried in the fold or in another interface. An NES assigned within a folded domain is therefore a false positive — the Φ-residues that would engage Xpo1 are precisely those packed into the hydrophobic core. This is a known failure mode (Kosugi *et al*., 2014), exemplified by the “false” NES of c-Abl (Hantschel *et al*., 2005). Accessibility can be judged from experimental structures, AlphaFold models, or biochemically by comparing Xpo1 interactions of the isolated peptide with that of the intact client candidate.

Finally, NES activity can be switched by covalent modification, for example at cysteine, which is preferred at most spacer positions and even at Φ^3^. An NES with cysteine at Φ^3^ should be inactivated by S-oxidation (to sulfenic, sulfinic, or sulfonic acid), S-nitrosylation, or disulfide formation, making it a sensor for these reactions. A serine or threonine at position 15 (Φ^4^+1) would likewise lose activity upon phosphorylation, where negative charge is detrimental; conversely, phosphorylating a serine at or adjacent to Φ^0^ could raise affinity, where negative charge is favored (Fig. 1B–C).

Prediction from sequence can be combined with experimental evidence for Xpo1 binding, and the two are complementary. Deep proteomics has mapped Xpo1 clients on a large scale — on the order of 1000 each in human and frog and ∼700 in yeast, including scores for the strength of the interaction (Kirli *et al*., 2015) with further human and frog datasets adding coverage (Thakar *et al*., 2013; Wühr *et al*., 2015). These datasets do not resolve whether a protein binds Xpo1 through its own NES or through a partner subunit, but they constrain the sequence-based prediction at the protein level: a high-scoring stretch in a documented Xpo1 client gains support, whereas the same stretch in a non-binding protein is probably a false positive — explicable, for instance, by burial or by an inactivating modification. Because both the preference-based scan and the interaction data are proteome-wide, their combination lifts NES assignment from the level of the individual peptide to that of the entire proteome.

Even with the constraints at the Φ- and spacer positions, Xpo1 accommodates an astronomical number of sequences that bind with modest affinity. We regard this degeneracy as a feature, because it makes NESs remarkably easy to acquire. Φ^0^ is dispensable, so only four Φ-positions must be matched; a longer segment offers many registers in which a match can occur; and leucine, a top-scoring residue at all Φ-positions, is the most abundant amino acid and is encoded by six of the 64 codons, so random coding sequence is already biased toward the required pattern. Above all, amphipathic helices — ubiquitous in α-helical proteins — display precisely the i, i+3/i+4 hydrophobic periodicity that an NES requires. Appending such a segment is therefore a plausible route to an export signal, and NESs are readily transplantable. This may explain why so many proteins acquired an NES during evolution — for biosynthetic transport, for keeping cytoplasmic factors out of nuclei, or for regulating cellular processes. It also explains why NES prediction is so error-prone: the pattern is common precisely because amphipathic helices are common, but in folded proteins their hydrophobic face is usually packed away (see above).

Within this vast space, however, there is also an optimum, and it is highly constrained — identical or close to hiNES4. Its affinity for Xpo1·RanGTP is low picomolar, comparable to that of exceptional antibodies or nanobodies, and 19000-fold higher than that of the wild-type PKI NES. Yet nature stops far short of it. As shown above, such affinities cause an irreversible arrest of the transport cycle — best explained by a failure to switch Xpo1 to its Ran-free, low-affinity state, from which the NES would normally be released. We propose that the GAP co-activators RanBP1/RanBP2, although they bind RanGTP tightly (K_D_ ∼0.1 nM), cannot extract the RanGTP·RanBP1/RanBP2 subcomplex from a hyperstabilized hiNES export complex and present it to RanGAP for GTPase activation. Such a complex would pose too high an energy barrier for catalytic disassembly.

Natural NESs therefore remain modest in affinity — not because tighter ones are out of reach (we built them). Xpo1 is a shared carrier that must serve a large molar excess of diverse cargoes, and this requires rapid transport cycles with fast release at the destination. A single super-high-affinity NES would sequester the exportin and, as our hiNESs demonstrate, stall export for all other cargoes *in trans*. The modest affinity of cellular NESs is thus not a per-cargo compromise but a global constraint on the entire NES repertoire of an organism.

## Materials and methods

### Library design, phage display, and deep sequencing

The NES library, representing wild type and all single-point mutations of the PKI NES, was generated by scanning a single NNK codon through the 15 NES codons. For this, 15 oligonucleotides with matching phagemid overhangs were synthesized, pooled, PCR-amplified, and cloned into a phagemid by circular polymerase extension cloning (Quan and Tian, 2011).

Handling of phages and selection followed (Pleiner *et al*., 2015). The selection strategy is described in the main text and Fig. 1A. Selection was performed in 50 mM Na·HEPES pH 7.4, 150 mM KCl, 10% glycerol, 0.1% BSA, 10 mM DTT. The 400-µl binding reaction contained 7×10^12^ phages, 2.7 µM Xpo1 (∼500-fold excess over phages) and 3.6 µM RanGTP and was incubated for 2 h; two 10-min washes with 4 ml selection buffer each then preceded protease elution.

The peptide-coding fragments of the phage DNA were amplified by PCR and used for library preparation with the TruSeq DNA PCR-Free Low Throughput Library Prep Kit (Illumina, Cat. No. 20015962); the clean-up step used 3:1 agarose (SERVA, Cat. No. 11385.02). The libraries were analyzed on a Fragment Analyzer and sequenced on a NextSeq 500 (150 cycles). Since the library was of low base diversity and flanked by constant regions, reliable base-calling required a 30% PhiX spike-in.

The sequencing data were analyzed with a custom script that filtered out low-quality reads (those with at least one base below Q13, corresponding to <95% base-call accuracy) or reads encoding a variant with more than one residue change. After filtering, 8M reads remained for the input (pre-selected) library, 1.2M for the background-binding library, 9M for the hXpo1-selected, and 10M for the yXpo1-selected library. The background was then subtracted from the selected-library reads, using a weighting factor that accounted for the ∼3000-fold lower phage recovery in the background than in the Xpo1 selections.

For each variant, enrichments were calculated as follows. First, its reads were counted in the input and in each selected library. Second, its relative abundance within each library was calculated by dividing its read count by the total number of reads in that library. Third, the enrichment upon Xpo1 selection was obtained by dividing the relative abundance in the selected library by that in the input library. Fourth, these enrichments were normalized to that of the wild-type PKI NES.

Finally, we had to account for the fact that the reference PKI NES sequence is not neutral: its residues are already near-optimal at some positions but suboptimal at others. Because all enrichments so far are expressed relative to this reference (step 4), an unusually strong or weak reference residue at a given position would distort the apparent preferences there. To correct for this, we applied a position-specific normalization, dividing all enrichments at that position by the same value — the ‘positional mutation enrichment’ (Fig. EV1A). This removed the position-specific bias.

Two further corrections were introduced. The first concerns the proline substitution at Φ² (position 9). Although clearly selected against during phage display, its value was higher than for proline at other positions. We therefore tested this substitution in transfection assays (Fig. EV2D) and found a complete loss of export activity, not only in a PKI context but even in a hiNES context. This justified setting the value to zero. Second, we noticed a codon-specific deviation for the alanine substitution at Φ³ (position 12): when encoded by GCG, it showed the expected strong negative selection, but as a GCT codon it appeared nearly neutral (possibly because of an amplification bias during sequencing). We resolved this discrepancy by transfection experiments, finding that a PKI NES carrying L12A has zero export activity (Fig. EV2D); we therefore based the preference value for this substitution on the GCG counts alone. Taken together, these steps yielded the preferences shown in Fig. 1B, C.

### Transient transfections of HeLa cells

Constructs for expressing NES fusions under the control of an eEF1A promoter were based on the pEGFP-C1 derivatives described in (Kirli *et al*., 2015). HeLa Kyoto cells were cultivated per ATCC guidelines, grown in 8-well slides (Ibidi, Cat. No. 80826), and co-transfected with the EGFP–NES construct of choice and an mCherry co-reporter — either mCherry fused to a wild-type PKI NES, or mCherry fused to both an SV40 NLS and a PKI NES — using Lipofectamine 3000 (Invitrogen, Cat. No. L3000001) per the manufacturer’s instructions.

Before imaging, the medium was replaced with phenol red-free Leibovitz’s L-15 medium containing 1.6 µM Hoechst 33342, and the slides were incubated under standard culturing conditions for 15 min.

Live cells were imaged on a TCS SP5 or SP8 confocal microscope (Leica Microsystems) with a 63×oil-immersion objective, using 405-, 488-, and 561-nm laser lines to excite Hoechst 33342, EGFP, and mCherry, respectively. Channels were acquired sequentially with laser settings adjusted per image to avoid saturation and keep the signal in the linear range. During imaging, samples were kept at 37 °C on a heated stage. If needed, images were linearly adjusted in the Fiji distribution of ImageJ2 (Schindelin *et al*., 2015).

For LMB treatment, the medium of cells (24 h post-transfection) was replaced with phenol red-free Leibovitz’s L-15 medium containing 10% FBS and 9 nM LMB (or the corresponding amount of ethanol, the solvent for the LMB stock), and cells were incubated under standard culturing conditions for 60 or 90 min before imaging.

### Recombinant protein expression and purification

Proteins used in this study were produced recombinantly (for the list of the expression constructs see Appendix Table S5) in *E. coli* cells of NEBExpress strain (New England Biolabs, Cat. No. C2523) and purified by immobilized metal ion affinity chromatography (IMAC) with elution either by protease elution (Frey and Görlich, 2014, 2015; Vera Rodriguez *et al*., 2019) or imidazole elution. As a polishing step, the proteins were purified by size exclusion chromatography (SEC) using Superdex 75 or Superdex 200 in prepacked HiLoad columns (Cytiva).

### FG phase assay

The assay compares, in a competitive setup, the affinities of two NESs (NES1 and NES2) for Xpo1 — or, to be precise, for the Xpo1·RanGTP subcomplex. Here, ‘competitive’ means that Xpo1 was kept well below the NES concentrations; equal concentrations of NES1 and NES2 were used for simplicity. The readout is the partitioning of the (spectrally separable) EGFP and mCherryR fusions into the Nup98 FG phase: the free NES fusion shows a partition coefficient of well below 1 (typically of ∼0.1; Figs. 2A, EV2A), whereas the NES·Xpo1·RanGTP complex reaches ≥1000 (Fig. 2A).

For each sample, a 50-µl analyte mix containing 3.75 µM NES-mCherryR, 3.75 µM NES-EGFP, 0.75 µM human Xpo1, and 3.75 µM hRanGTP(5–180, Q69L) in assay buffer (50 mM Tris/HCl pH 7.4, 150 mM NaCl, 5 mM MgCl₂, 100 µM TCEP) was prepared. The order of addition is critical here, because once formed, the NES·Xpo1·RanGTP complexes are kinetically very stable — with hiNES off-rates of at least days, and hence correspondingly long relaxation times. To prevent the assay from reporting ratios of on-rates rather than affinities, we first pre-mixed the NES fusions, then added the exportin and incubated for 1 h to allow an initial equilibrium to be established. Only then was RanGTP added, which essentially locks the equilibria in place.

The FG phase suspension (Schmidt and Görlich, 2015; Ng *et al*., 2021) was prepared by rapid 50-fold dilution of a 1 mM stock (in 4 M GdnHCl) of His-tagged MacNup98 GLFG domain (residues 1–696) from Tetrahymena thermophila into assay buffer, followed by further dilution to 7.5 µM.

25 µl of the FG phase suspension was mixed with 50 µl of the analyte mix and pipetted into the wells of an 18-well flat µ-slide (Ibidi, Cat. No. 81817) passivated with fish gelatin. The samples were incubated at room temperature in the dark for 20 min to allow the FG particles to settle, then covered with 50 µl paraffin oil to prevent evaporation. The sealed samples were then incubated for a further 40 min to allow the partitioning equilibrium to be established.

Images were collected on a Stellaris 5 confocal microscope (Leica Microsystems), equipped with a white-light laser and HyD S detectors, using a 63× water-immersion objective and sequential excitation of EGFP (488 nm; detection 500-550 nm) and mCherryR (587 nm; detection 600-650 nm). Minus EGFP and minus mCherryR controls confirmed that channel cross-talk, autofluorescence, and instrumental background were neglible for these settings and the concentrations of fluorescent proteins used.

Scanning was performed in Z-stacks (voxel size 120×120×300 nm) in photon-counting mode within the linear range of the detectors for precise quantification. Laser intensities (typically 0.1-0.5%) were selected to exploit the linear range of the detectors (about 400 photons per pixel for the brightes sample at 2.8 µs pixel dwell time with 2x line accumulation) and kept constant within an experiment unless stated otherwise.

For calculating partition coefficients, a median filter was applied to the 3D images, and thresholding was used to segment the FG particles. The gray values inside the particles were averaged (outer layer of 1.25 µm omitted, particles under 2.5 µm in diameter excluded from analysis), as were gray values in the adjacent buffer (excluding a 2-µm zone around the FG phase). The partition coefficient was calculated as the ratio of the average gray value in the phase to that in the buffer (typically 50-100 µm^3^ of the phase volume from 20-35 particles and 1000-2000 µm^3^ of buffer volume included in the analysis). Images were processed for presentation and analysis using the Fiji distribution of ImageJ2 (Schindelin *et al*., 2015) and the scikit-image Python library (van der Walt *et al*., 2014), respectively. Partition coefficients were used to calculate relative affinities for pairs of NESs; the derivation is provided in the Appendix Supplementary Methods. All the experiments were repeated at least twice with at least three 3D images collected per sample in each experiment.

### Anisotropy measurements

Maleimide labeling with fluorescein was done as described in (Pleiner *et al*., 2015). The assay buffer contained 20 mM HEPES/ NaOH pH 7.2, 150 mM KCl, 5 mM MgCl_2_, 0.01% BSA, 0.5 mM TCEP, and 50 µM GTP.

For the direct-binding measurements, 10 nM fluorescein-labeled NES-ZZ fusion, 2.5 µM RanGTP (where appropriate), and varying concentrations of Xpo1 up to 1 µM were used. For the competition experiments, 20 nM fluorescein-labeled NES-ZZ or S1 NES-ZZ fusion, varying concentrations of NES-MBP fusions, 50 nM or 30 nM Xpo1 and 1 µM RanGTP were used and added in this order. The proteins were mixed in transparent non-binding 96-well plates (Greiner Bio-One, Cat. No. 655901) and incubated for 4 h at room temperature in the dark.

Fluorescence anisotropy (Dandliker and Feigen, 1961; Güttler *et al*., 2010) was measured in black non-binding 96-well plates (Greiner Bio-One, Cat. No. 655900) using a Synergy H4 microplate reader (excitation 485/20 nm, emission 528/20 nm, sensitivity 65) at 23°C. The data were fitted to appropriate models given by the equation systems (1) and (2) below for anisotropy change in direct (Heyduk and Lee, 1990) or competitive binding (Wang, 1995) respectively using the LMFIT Python library (Newville, M. *et al*., 2014) to estimate K_D_s.

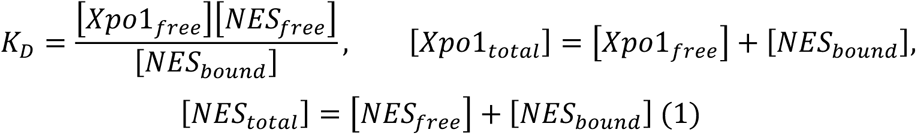

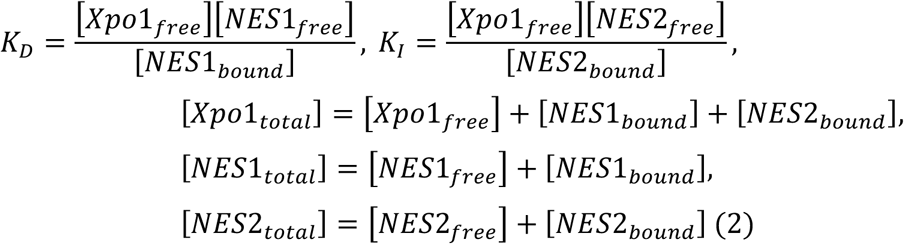

### Complex preparation and crystallization

yXpo1 (Δ377–413; Koyama and Matsuura, 2010) and yRan(1–182)(Q71L)·GTP were purified by IMAC with protease elution and SEC. For complex formation, H14-MBP-SUMOEu-hiNES2 was immobilized on IMAC resin and supplemented with an equimolar amount of yXpo1(Δ377–413) and a 1.4-fold excess of yRan(1–182)(Q71L)·GTP. Unbound proteins were washed away, and the complex was eluted with SENP^Eu^ protease (Vera Rodriguez *et al*., 2019). The complex was further purified by SEC in 20 mM Na·HEPES pH 7.4, 50 mM NaCl, and 1 mM TCEP.

The crystal used for structure determination was obtained at 20°C by a hanging-drop setup with 500 µl of reservoir solution (100 mM Tris/HCl pH 7.4, 12% PEG 20000, 5 mM Mg(OAc)₂) per well, by microseeding into drops containing 1.5 µl of 8.7 mg/ml complex solution, 1 µl of reservoir solution, and 0.5 µl of seed stock from crystals grown previously under similar conditions. Before data collection, the crystal was cryoprotected in 100 mM Tris/HCl pH 7.4, 12.5% PEG 20000, 5 mM Mg(OAc)₂, 20% ethylene glycol and snap-frozen in liquid nitrogen.

### Structure determination and analysis

Diffraction data were collected at beamline X10SA of the Swiss Light Source, Paul Scherrer Institute (Villigen, Switzerland), using a PILATUS 6M detector. Data were processed with DIALS (Beilsten-Edmands *et al*., 2020). The structure was solved by molecular replacement with Phaser (McCoy et al., 2007), using the Xpo1·PKI·Gsp1p·GTP complex (PDB 3WYG; Koyama *et al*., 2014) without the PKI chain as a search model. hiNES2 chains were built manually in Coot (Emsley *et al*., 2010). The structure was refined through alternating cycles of reciprocal-space refinement in REFMAC5 (Vagin *et al*., 2004), real-space refinement in Phenix (Liebschner et al., 2019), and manual rebuilding in Coot.

The asymmetric unit contained four complexes with only minor differences between them (Appendix Fig. S1A). All the figures are made with the complex with the best electron density (chains A, C, and I in the PDB) unless stated otherwise. In all four copies of the complex, hiNES2 adopted a nearly identical conformation (Appendix Fig. S1B) with the differences (RMSD of 0.8 Å to 1.1 Å between the best resolved reference chain and the other copies) mostly confined to the side chain rotamers, particularly for the solvent-exposed residues. The peptide could be unambiguously assigned to the electron density (Appendix Fig. S1C).

Interface analysis was performed with PISA (Krissinel and Henrick, 2007); the omit map was generated in Phenix. Conservation analysis was performed with UCSF Chimera (Pettersen *et al*., 2004). Figures were prepared with PyMOL (Schrödinger, LLC).

## Data availability

Coordinates and structure factors have been deposited with the Protein Data Bank (PDB) with the accession code 8QYZ.

## Acknowledgements

We thank Kerstin Maier and Dr. Kristina Žumer (Department of Molecular Biology, MPI NAT) for help with Illumina sequencing, Ulrich Steuerwald and Jürgen Wawrzinek (Crystallization Facility, MPI NAT) for help with the crystallization conditions screening and crystal freezing and shipment, the team of the SLS Beamlines (Paul Scherrer Institut, Switzerland) for help with the X-ray diffraction data collection, Dr. Sheung Chun Ng for providing the Nup98 FG domain, Susanne Brandfass for help with cloning and protein expression, Fatma Chafra, Jennifer Miao, and Aastha Vartak for critical reading of the manuscript. OR was supported by the Boehringer Ingelheim Fonds PhD fellowship.

## Expanded View figure legends

**Figure EV1.**
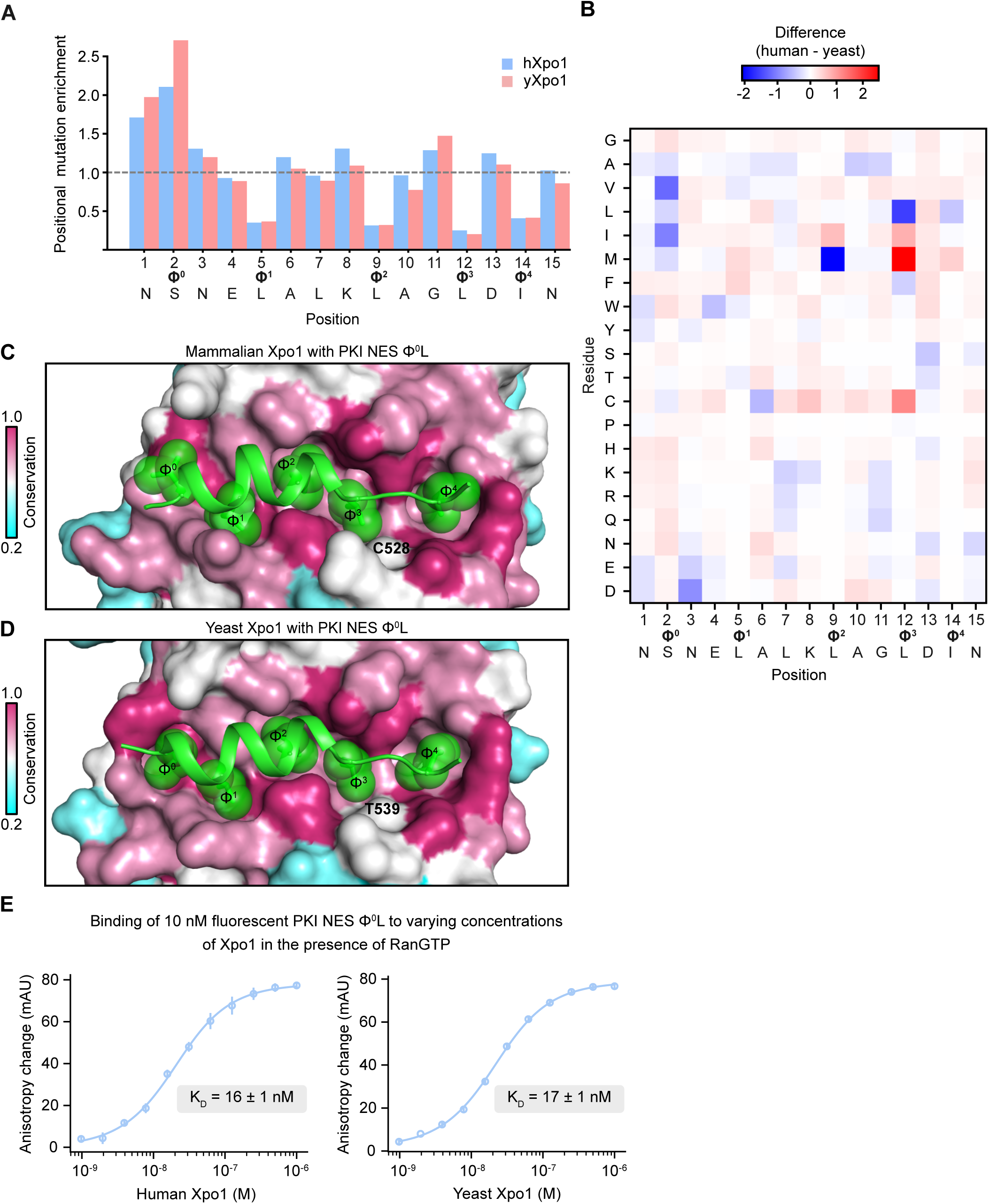
Comparison of human and yeast Xpo1: NES preferences, conservation, and affinity. **(A)** Positional mutation enrichment: the selected-to-naïve frequency ratio of mutations (to any non-WT residue) at each PKI NES position. A higher ratio means a selection against the original residue and that mutations at that position are enriched overall, regardless of the substituting residue. **(B)** Difference between the human (Fig. 1B) and yeast (Fig. 1C) preference heatmaps. Positive values (red) mark exchanges more favored by human Xpo1; negative values (blue), those more favored by yeast Xpo1. Asterisks mark two positions treated separately in the analysis (black, position 12; green, position 9; see Methods). (**C**, **D**) Surface representations of the NES-binding pocket of mammalian Xpo1 (**C**; *M. musculus*, PDB 3NBY, Güttler et al., 2010; nearly identical to human Xpo1) and fungal Xpo1 (**D**; *S. cerevisiae*, PDB 3WYG, Koyama et al., 2014), colored by residue conservation across five distant eukaryotic lineages (*H. sapiens*, *S. cerevisiae*, *T. thermophila*, *D. discoideum*, and *A. thaliana*). The PKI NES is green, with the Φ-residue side chains (and Cα) as sticks and semi-transparent spheres, labeled. C528/T539 (bold) are the least conserved residues near the bound NES. (**E**) Fluorescence anisotropy of 10 nM fluorescently labeled PKI NES SΦ⁰L-ZZ titrated with hXpo1 and excess hRanGTP(5–180, Q69L) (left) or yXpo1 and yRanGTP(1–182, Q71L) (right). K_D_s were obtained by fitting a direct-binding model (Heyduk and Lee, 1990). Error bars, standard error of technical triplicates.

**Figure EV2.**
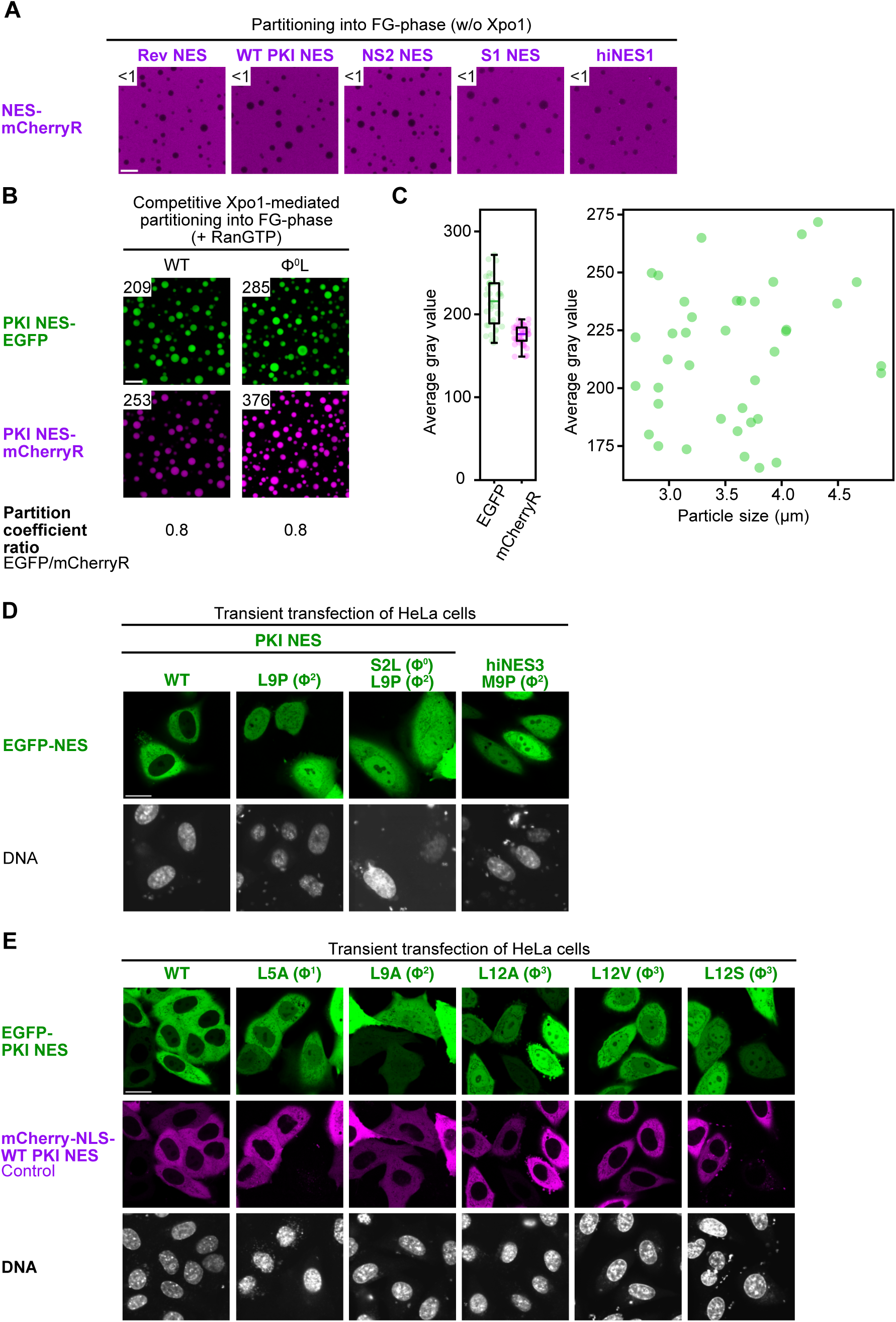
Controls for the FG phase assay and additional NES mutants. **(A)** In the absence of Xpo1, all NES-mCherryR fusions used in this study are excluded from the FG phase (partition coefficient <1), confirming that their partitioning in the main assay is Xpo1-dependent. Scale bar, 10 µm. **(B)** On the left, WT PKI NES competed as an EGFP fusion against an mCherryR fusion for limiting Xpo1 in an FG partitioning assay. The right shows the same for the Ö⁰L variant. For both NESs, the EGFP:mCherryR partition-coefficient ratio is the same (∼0.8. Thus, the two tags still carry a slight bias, but a constant one that will cancel out upon normalization. Scale bar, 10 µm. **(C)** Distribution of per-particle signal (left) and its relationship to particle size (right) in the EGFP channel, for the WT PKI NES sample in (B). r, Pearson correlation coefficient. **(D)** Proline-scan mutants — L9P (Φ²) and S2L(Φ⁰) L9P(Φ²) of the PKI NES, and M9P(Φ²) of hiNES3 — fused to the C-terminus of EGFP and expressed in cells. In the phage selection, L9P was selected against but (apparently artifactually) less strongly than expected for a helix-breaking proline at Φ². The cell-based assay shown here confirms that the proline abolishes export even in the strongest context (a hiNES). Scale bar, 20 µm. **(E)** Indicated PKI NES variants, fused to EGFP, were assayed in cells for export activity. Leucine-to-alanine substitutions at Ö¹, Φ², and Φ³ were detrimental, with a possibly residual activity for L5A (Φ¹), but a complete loss for L9A and L12A. Further Φ³ substitutions (L12V, L12S) were included as controls and were likewise inactive. That even the conservative L12V abolishes export is notable, but consistent with its depletion in the phage selection. Scale bar, 20 µm.

## Appendix

### Appendix Tables

**Appendix Table S1.** Positional preferences of human Xpo1 as represented by the heatmap of Fig. 1B.

|  | 1 | 2 | 3 | 4 | 5 | 6 | 7 | 8 | 9 | 10 | 11 | 12 | 13 | 14 | 15 |
| --- | --- | --- | --- | --- | --- | --- | --- | --- | --- | --- | --- | --- | --- | --- | --- |
| G | 1.009 | 1.141 | 1.029 | 0.854 | 0.076 | 0.608 | 0.620 | 0.496 | 0.017 | 2.268 | 1.074 | 0.109 | 0.566 | 0.023 | 1.597 |
| A | 1.263 | 1.357 | 1.068 | 0.606 | 0.107 | 1.153 | 1.242 | 0.686 | 0.120 | 1.436 | 1.149 | 0.100 | 0.855 | 0.018 | 1.240 |
| V | 1.170 | 2.335 | 1.348 | 1.021 | 1.136 | 1.209 | 1.376 | 0.575 | 0.711 | 0.332 | 0.881 | 0.608 | 1.012 | 0.791 | 0.558 |
| L | 1.418 | 2.067 | 1.602 | 0.869 | 3.935 | 1.480 | 1.442 | 2.017 | 4.402 | 0.565 | 0.902 | 5.514 | 1.063 | 6.557 | 0.398 |
| I | 1.549 | 2.862 | 1.243 | 0.962 | 2.806 | 1.522 | 1.777 | 0.965 | 2.426 | 0.649 | 0.674 | 2.397 | 0.954 | 3.399 | 0.490 |
| M | 1.348 | 2.109 | 1.405 | 0.999 | 1.244 | 0.990 | 1.092 | 1.324 | 7.538 | 0.478 | 0.895 | 7.937 | 1.306 | 2.444 | 0.419 |
| F | 1.328 | 1.220 | 0.986 | 0.830 | 1.249 | 0.494 | 1.086 | 0.928 | 5.974 | 0.536 | 0.626 | 2.000 | 1.326 | 0.173 | 0.500 |
| W | 1.501 | 1.197 | 1.997 | 0.986 | 0.203 | 0.946 | 0.746 | 0.955 | 0.157 | 0.417 | 0.845 | 0.141 | 0.714 | 0.170 | 0.551 |
| Y | 1.150 | 1.179 | 1.478 | 0.794 | 0.082 | 0.704 | 0.839 | 0.585 | 0.176 | 0.354 | 0.335 | 0.064 | 0.922 | 0.080 | 0.708 |
| S | 1.048 | 0.656 | 0.818 | 0.791 | 0.064 | 1.245 | 1.239 | 0.819 | 0.033 | 1.773 | 0.744 | 0.050 | 1.558 | 0.011 | 1.706 |
| T | 1.115 | 1.158 | 1.167 | 0.934 | 0.439 | 1.389 | 0.859 | 0.552 | 0.263 | 0.293 | 0.627 | 0.031 | 0.834 | 0.108 | 1.334 |
| C | 1.049 | 1.782 | 1.541 | 1.483 | 0.777 | 2.041 | 1.893 | 1.902 | 0.329 | 1.538 | 1.733 | 2.237 | 1.645 | 0.052 | 2.171 |
| P | 0.726 | 1.027 | 0.564 | 0.426 | 0.110 | 0.303 | 0.080 | 0.125 | 0.000 | 0.089 | 0.021 | 0.100 | 0.040 | 0.047 | 0.054 |
| H | 0.998 | 0.852 | 0.848 | 0.638 | 0.065 | 0.851 | 0.948 | 0.565 | 0.114 | 0.718 | 1.021 | 0.008 | 1.343 | 0.004 | 1.544 |
| K | 0.865 | 0.476 | 0.861 | 0.463 | 0.114 | 0.049 | 1.146 | 1.056 | 0.058 | 1.139 | 1.576 | 0.045 | 1.077 | 0.017 | 1.382 |
| R | 0.835 | 0.378 | 0.404 | 0.258 | 0.037 | 0.229 | 0.834 | 0.997 | 0.033 | 0.702 | 2.139 | 0.020 | 0.662 | 0.038 | 1.405 |
| Q | 1.077 | 0.869 | 1.085 | 0.964 | 0.047 | 0.570 | 1.279 | 0.754 | 0.118 | 1.077 | 1.200 | 0.083 | 1.281 | 0.007 | 1.088 |
| N | 0.808 | 0.955 | 1.057 | 0.882 | 0.083 | 0.906 | 1.499 | 0.250 | 0.063 | 0.597 | 0.727 | 0.016 | 1.390 | 0.176 | 1.346 |
| E | 1.177 | 1.207 | 1.464 | 1.491 | 0.128 | 1.051 | 1.509 | 0.540 | 0.068 | 0.818 | 1.102 | 0.108 | 1.239 | 0.012 | 0.204 |
| D | 1.366 | 1.563 | 1.905 | 1.874 | 0.075 | 0.596 | 1.418 | 0.423 | 0.014 | 1.224 | 1.454 | 0.069 | 1.107 | 0.019 | 0.534 |

**Appendix Table S2.** Positional preferences of yeast Xpo1 as represented by the heatmap of Fig. 1C.

|  | 1 | 2 | 3 | 4 | 5 | 6 | 7 | 8 | 9 | 10 | 11 | 12 | 13 | 14 | 15 |
| --- | --- | --- | --- | --- | --- | --- | --- | --- | --- | --- | --- | --- | --- | --- | --- |
| G | 0.928 | 0.819 | 0.890 | 0.747 | 0.072 | 0.547 | 0.480 | 0.395 | 0.014 | 2.047 | 0.968 | 0.147 | 0.359 | 0.025 | 1.546 |
| A | 1.411 | 1.613 | 1.060 | 0.658 | 0.196 | 1.361 | 1.437 | 0.677 | 0.098 | 1.843 | 1.491 | 0.103 | 0.705 | 0.012 | 1.135 |
| V | 1.183 | 3.546 | 1.171 | 0.903 | 1.308 | 1.259 | 1.429 | 0.457 | 0.493 | 0.295 | 0.623 | 0.457 | 0.834 | 0.595 | 0.492 |
| L | 1.442 | 2.391 | 1.386 | 0.851 | 3.897 | 1.169 | 1.599 | 1.941 | 4.447 | 0.593 | 1.014 | 7.077 | 0.732 | 7.023 | 0.401 |
| I | 1.643 | 3.891 | 1.125 | 0.841 | 2.657 | 1.424 | 1.879 | 0.707 | 1.769 | 0.681 | 0.486 | 1.584 | 0.724 | 3.441 | 0.503 |
| M | 1.419 | 2.389 | 1.366 | 1.077 | 0.862 | 0.792 | 1.139 | 1.204 | 9.611 | 0.545 | 0.967 | 5.335 | 1.052 | 1.973 | 0.416 |
| F | 1.158 | 1.063 | 0.895 | 0.713 | 0.839 | 0.403 | 0.913 | 0.911 | 5.828 | 0.450 | 0.465 | 2.387 | 1.009 | 0.163 | 0.411 |
| W | 1.772 | 0.874 | 1.857 | 1.496 | 0.343 | 0.931 | 0.681 | 0.701 | 0.185 | 0.464 | 0.678 | 0.176 | 0.385 | 0.182 | 0.449 |
| Y | 1.330 | 1.142 | 1.618 | 0.733 | 0.087 | 0.615 | 0.814 | 0.668 | 0.109 | 0.352 | 0.260 | 0.068 | 0.788 | 0.038 | 0.697 |
| S | 1.026 | 0.527 | 0.803 | 0.759 | 0.048 | 1.163 | 1.172 | 0.628 | 0.034 | 1.782 | 0.755 | 0.055 | 2.010 | 0.009 | 1.861 |
| T | 1.064 | 1.197 | 1.135 | 0.853 | 0.566 | 1.116 | 0.897 | 0.378 | 0.136 | 0.322 | 0.511 | 0.035 | 1.075 | 0.079 | 1.284 |
| C | 0.928 | 1.940 | 1.396 | 1.185 | 0.763 | 2.611 | 1.588 | 1.327 | 0.149 | 1.206 | 1.533 | 1.030 | 1.725 | 0.028 | 1.958 |
| P | 0.767 | 0.940 | 0.464 | 0.399 | 0.103 | 0.215 | 0.066 | 0.133 | 0.000 | 0.101 | 0.012 | 0.134 | 0.040 | 0.038 | 0.049 |
| H | 0.779 | 0.575 | 0.865 | 0.557 | 0.063 | 0.549 | 0.976 | 0.486 | 0.114 | 0.647 | 0.924 | 0.007 | 1.504 | 0.007 | 1.363 |
| K | 0.696 | 0.264 | 0.844 | 0.439 | 0.088 | 0.046 | 1.518 | 1.311 | 0.055 | 1.012 | 1.790 | 0.050 | 0.977 | 0.009 | 1.110 |
| R | 0.700 | 0.202 | 0.448 | 0.259 | 0.040 | 0.150 | 1.101 | 1.036 | 0.030 | 0.750 | 2.355 | 0.028 | 0.575 | 0.031 | 1.231 |
| Q | 1.020 | 0.603 | 1.106 | 0.904 | 0.051 | 0.475 | 1.473 | 0.737 | 0.117 | 1.128 | 1.520 | 0.101 | 1.217 | 0.008 | 1.085 |
| N | 0.722 | 0.640 | 1.191 | 0.792 | 0.077 | 0.550 | 1.394 | 0.230 | 0.047 | 0.525 | 0.748 | 0.020 | 1.665 | 0.178 | 1.662 |
| E | 1.416 | 1.116 | 1.841 | 1.606 | 0.127 | 0.858 | 1.760 | 0.459 | 0.057 | 0.700 | 1.216 | 0.149 | 1.070 | 0.011 | 0.279 |
| D | 1.600 | 1.444 | 2.826 | 1.951 | 0.074 | 0.613 | 1.197 | 0.378 | 0.009 | 0.897 | 1.248 | 0.081 | 1.295 | 0.010 | 0.625 |

**Appendix Table S3.** Scoring Matrix used for computing the prediction scores in the Table 1.

|  | 1 | 2 | 3 | 4 | 5 | 6 | 7 | 8 | 9 | 10 | 11 | 12 | 13 | 14 | 15 |
| --- | --- | --- | --- | --- | --- | --- | --- | --- | --- | --- | --- | --- | --- | --- | --- |
| G | 1.249 | 1.740 | 0.973 | 0.573 | 0.019 | 0.527 | 0.430 | 0.470 | 0.004 | 1.579 | 1.000 | 0.020 | 0.512 | 0.007 | 1.186 |
| A | 1.564 | 2.069 | 1.010 | 0.407 | 0.027 | 1.000 | 0.861 | 0.649 | 0.027 | 1.000 | 1.070 | 0.018 | 0.773 | 0.005 | 0.921 |
| V | 1.448 | 3.561 | 1.275 | 0.685 | 0.289 | 1.049 | 0.954 | 0.544 | 0.162 | 0.231 | 0.820 | 0.110 | 0.915 | 0.233 | 0.415 |
| L | 1.756 | 3.152 | 1.515 | 0.583 | 1.000 | 1.283 | 1.000 | 1.909 | 1.000 | 0.394 | 0.840 | 1.000 | 0.960 | 1.929 | 0.296 |
| I | 1.918 | 4.364 | 1.176 | 0.645 | 0.713 | 1.319 | 1.232 | 0.914 | 0.551 | 0.452 | 0.627 | 0.435 | 0.862 | 1.000 | 0.364 |
| M | 1.669 | 3.216 | 1.328 | 0.670 | 0.316 | 0.858 | 0.758 | 1.254 | 1.712 | 0.333 | 0.834 | 1.439 | 1.180 | 0.719 | 0.311 |
| F | 1.644 | 1.860 | 0.932 | 0.557 | 0.317 | 0.428 | 0.753 | 0.879 | 1.357 | 0.373 | 0.583 | 0.363 | 1.198 | 0.051 | 0.371 |
| W | 1.859 | 1.826 | 1.889 | 0.661 | 0.051 | 0.821 | 0.517 | 0.904 | 0.036 | 0.290 | 0.787 | 0.026 | 0.645 | 0.050 | 0.409 |
| Y | 1.425 | 1.798 | 1.398 | 0.533 | 0.000 | 0.610 | 0.582 | 0.554 | 0.040 | 0.247 | 0.312 | 0.000 | 0.833 | 0.000 | 0.526 |
| S | 1.298 | 1.000 | 0.774 | 0.531 | 0.000 | 1.079 | 0.859 | 0.776 | 0.000 | 1.234 | 0.693 | 0.000 | 1.407 | 0.000 | 1.267 |
| T | 1.380 | 1.765 | 1.104 | 0.626 | 0.112 | 1.204 | 0.596 | 0.523 | 0.060 | 0.204 | 0.584 | 0.006 | 0.754 | 0.032 | 0.991 |
| C | 1.299 | 2.718 | 1.458 | 0.995 | 0.198 | 1.769 | 1.313 | 1.801 | 0.075 | 1.070 | 1.613 | 0.406 | 1.486 | 0.015 | 1.612 |
| P | 0.899 | 1.566 | 0.533 | 0.286 | 0.000 | 0.262 | 0.056 | 0.118 | 0.000 | 0.062 | 0.020 | 0.000 | 0.036 | 0.000 | 0.040 |
| H | 1.236 | 1.300 | 0.802 | 0.428 | 0.000 | 0.738 | 0.658 | 0.535 | 0.000 | 0.500 | 0.951 | 0.000 | 1.213 | 0.000 | 1.147 |
| K | 1.071 | 0.727 | 0.814 | 0.311 | 0.000 | 0.042 | 0.795 | 1.000 | 0.000 | 0.793 | 1.468 | 0.000 | 0.973 | 0.000 | 1.026 |
| R | 1.034 | 0.576 | 0.382 | 0.173 | 0.000 | 0.199 | 0.578 | 0.944 | 0.000 | 0.489 | 1.991 | 0.000 | 0.598 | 0.000 | 1.043 |
| Q | 1.334 | 1.324 | 1.026 | 0.647 | 0.000 | 0.494 | 0.887 | 0.714 | 0.000 | 0.750 | 1.117 | 0.000 | 1.157 | 0.000 | 0.808 |
| N | 1.000 | 1.457 | 1.000 | 0.592 | 0.000 | 0.785 | 1.040 | 0.236 | 0.000 | 0.416 | 0.677 | 0.000 | 1.256 | 0.000 | 1.000 |
| E | 1.458 | 1.840 | 1.384 | 1.000 | 0.000 | 0.912 | 1.047 | 0.511 | 0.000 | 0.570 | 1.026 | 0.000 | 1.120 | 0.000 | 0.152 |
| D | 1.691 | 2.383 | 1.802 | 1.257 | 0.000 | 0.517 | 0.983 | 0.400 | 0.000 | 0.852 | 1.354 | 0.000 | 1.000 | 0.000 | 0.397 |

**Appendix Table S4.** Crystallographic data collection and refinement statistics for the hiNES2·yXpo1·RanGTP structure (PDB ID: 8QYZ).

|  |  |
| --- | --- |
| **Space group** | P1 |
| **Data collection** |  |
| Unit cell dimensions |  |
| a, b, c (Å) | 97.4, 105.6, 170.4 |
| α, β, γ (°) | 82.1, 86.7, 76.7 |
| Multiplicity | 3.41 (3.25)* |
| Completeness (%) | 96.34 (95.73) |
| R<sub>pim</sub> | 0.090 (0.656) |
| R<sub>merge</sub> | 0.1386 (0.9952) |
| I/σI | 8.05 (0.83) |
| CC1/2 | 0.8256 (0.6060) |
| **Refinement** |  |
| Resolution range | 59.96–3.0 (3.107–3.0) |
| No. of reflections | 126220 (12521) |
| R<sub>work</sub> | 0.2177 (0.3191) |
| R<sub>free</sub> | 0.2470 (0.3402) |
| No. of atoms |  |
| Protein | 38909 |
| Ligands | 148 |
| Solvent | 209 |
| B-factors (Å<sup>2</sup>) |  |
| Protein | 80.09 |
| Ligands | 83.17 |
| Solvent | 38.62 |
| RMSDs |  |
| Bonds (Å) | 0.007 |
| Angles (°) | 1.35 |
| Ramachandran analysis |  |
| Favored (%) | 98.06 |
| Allowed (%) | 1.94 |
| Outliers (%) | 0 |
| Rotamer outliers (%) | 1.90 |
| Clashscore | 9.21 |
| No. of TLS groups | 12 |

\* Statistics for the highest-resolution shell are shown in parentheses.

**Appendix Table S5.**
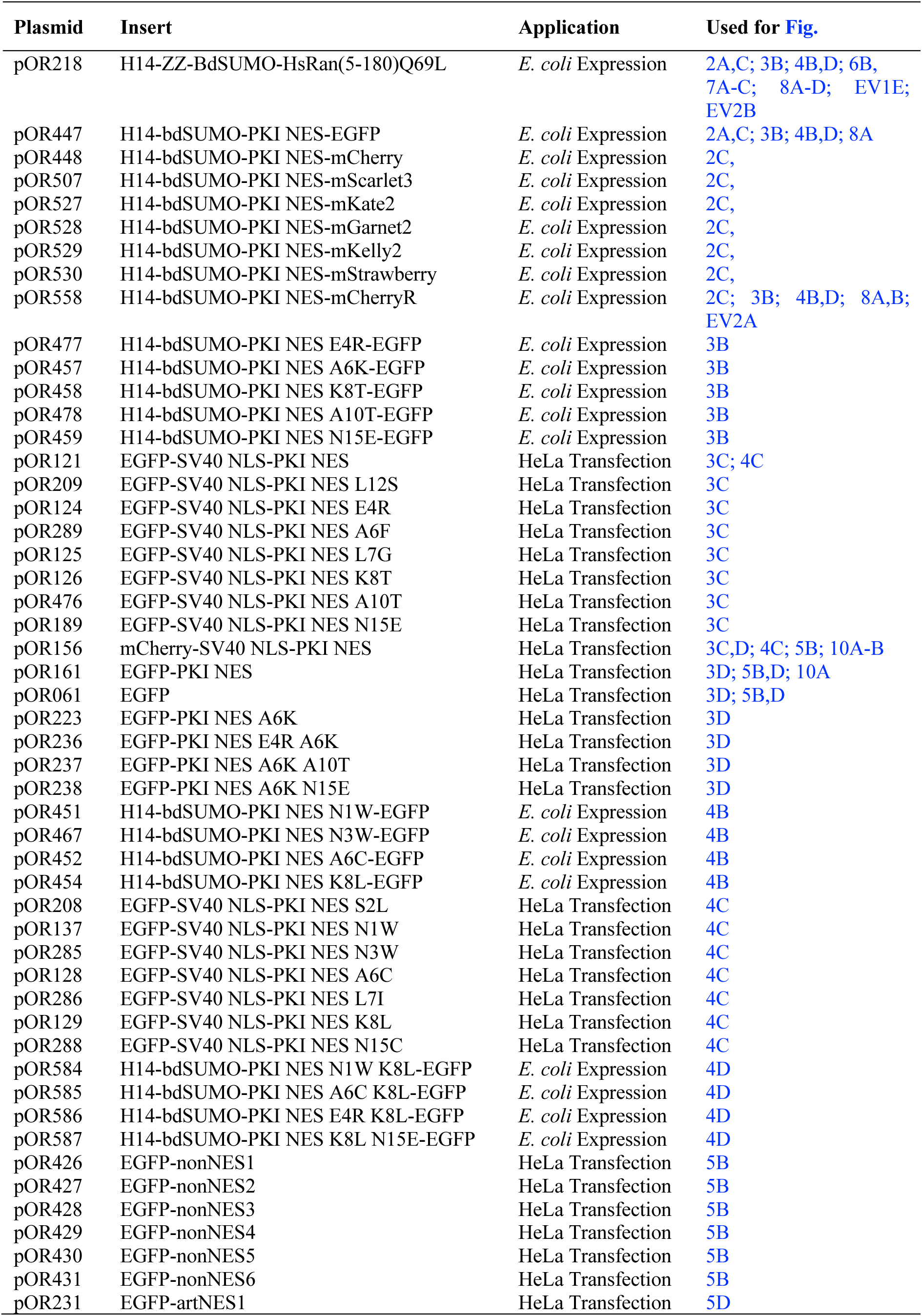
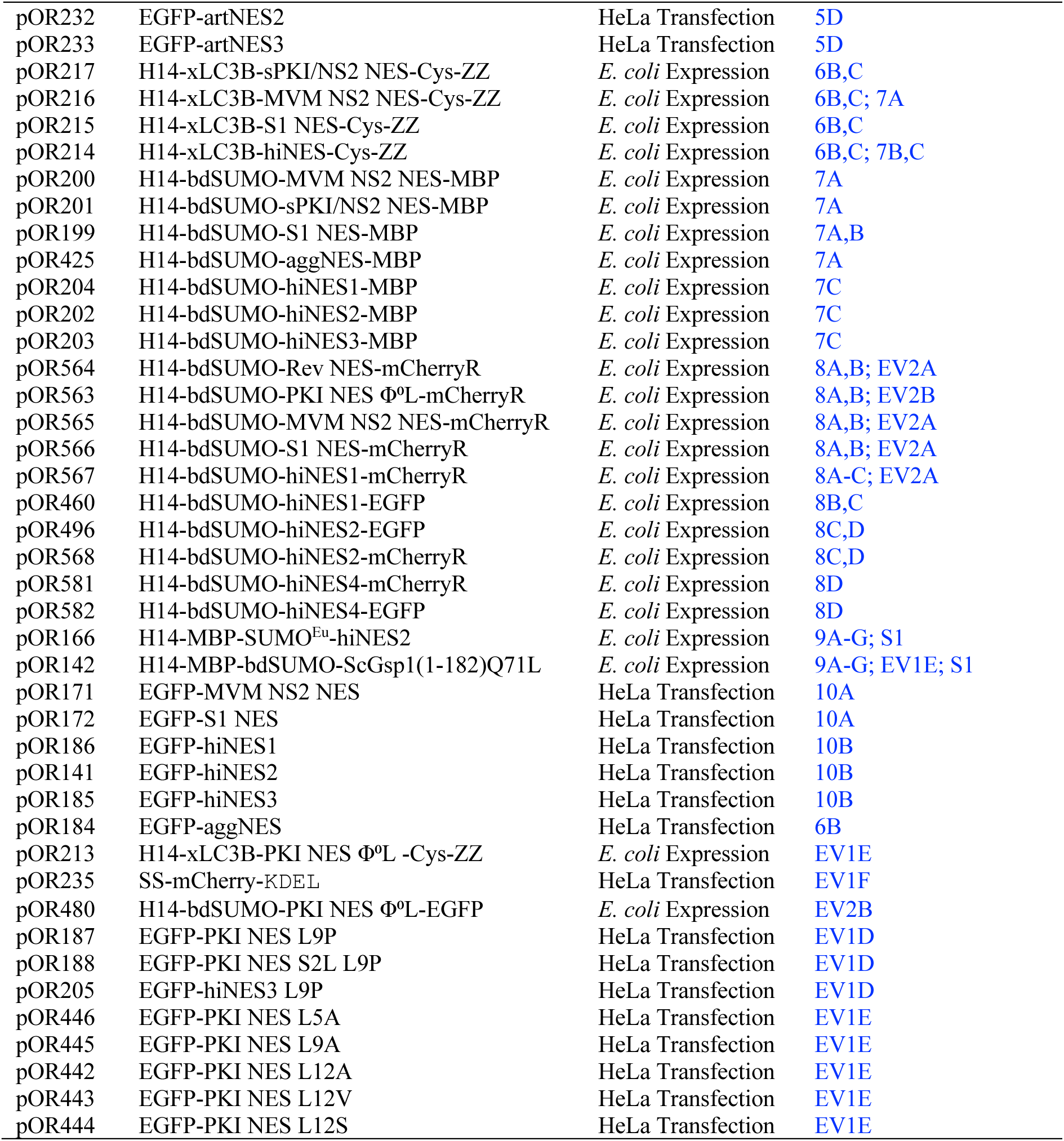
Plasmids created and used in this study.

### Appendix Figure

**Appendix Figure S1.**
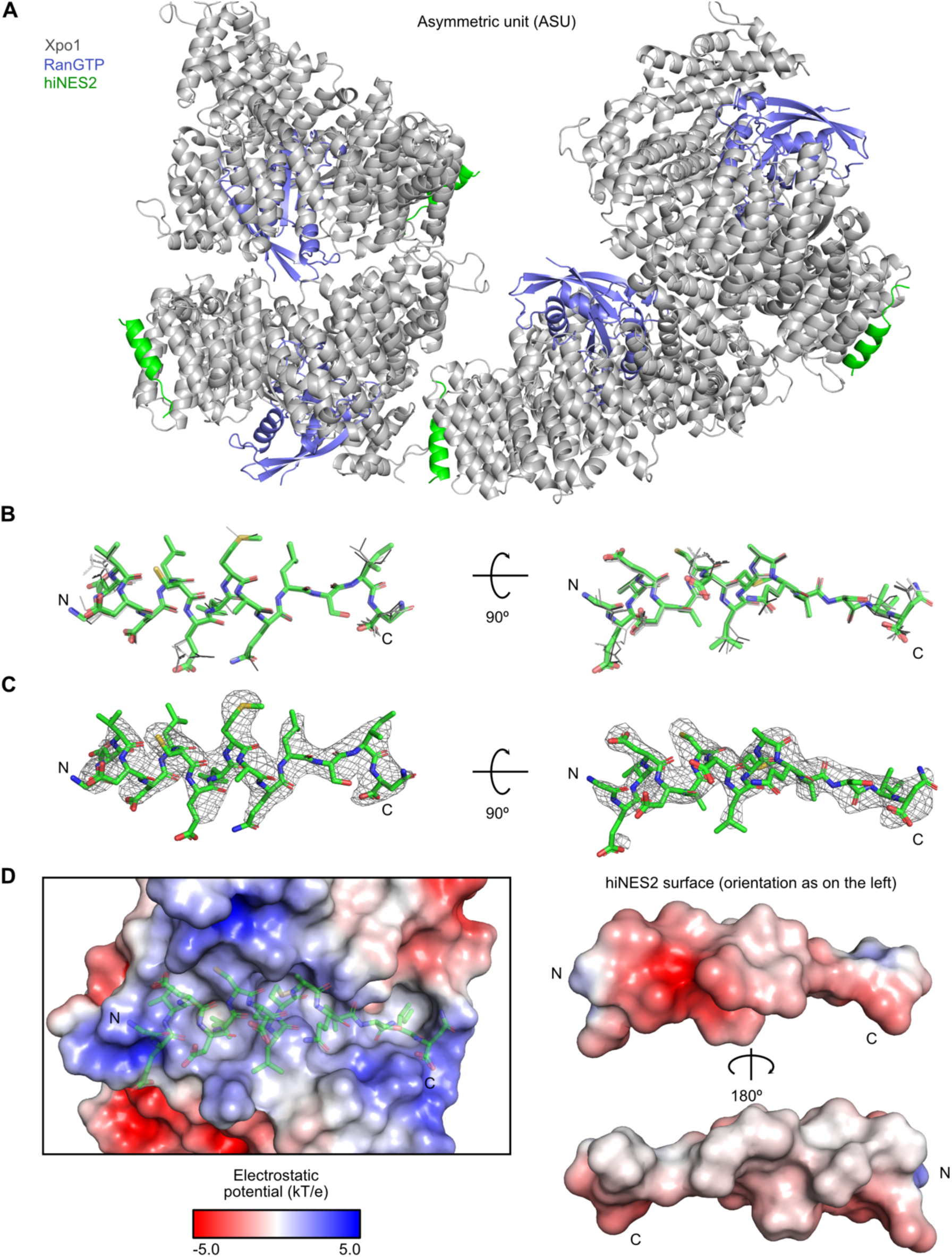
Non-crystallographic symmetry, hiNES2 model quality, and charge complementarity in the hiNES2·yXpo1·RanGTP complex. **(A)** Asymmetric unit of the crystal, containing four copies of the complex in virtually the same relative arrangement. **(B)** Superposition of the four hiNES2 copies in the asymmetric unit. The best-resolved chain is shown as green sticks; the others as thinner gray sticks. **(C)** hiNES2 (green sticks) with the omit map (all hiNES2 chains omitted) contoured at 3.0σ. **(D)** Charge complementarity between the yXpo1 peptide-binding pocket and hiNES2. yXpo1 and hiNES2 (two orientations) are shown as surfaces colored by electrostatic potential (kT/e). hiNES2 is also shown in the binding pocket as semi-transparent green sticks. The hiNES2 termini (N, C) are labeled.

### Appendix Supplementary Material

#### Sequences of the NES fusions for the FG phase assay

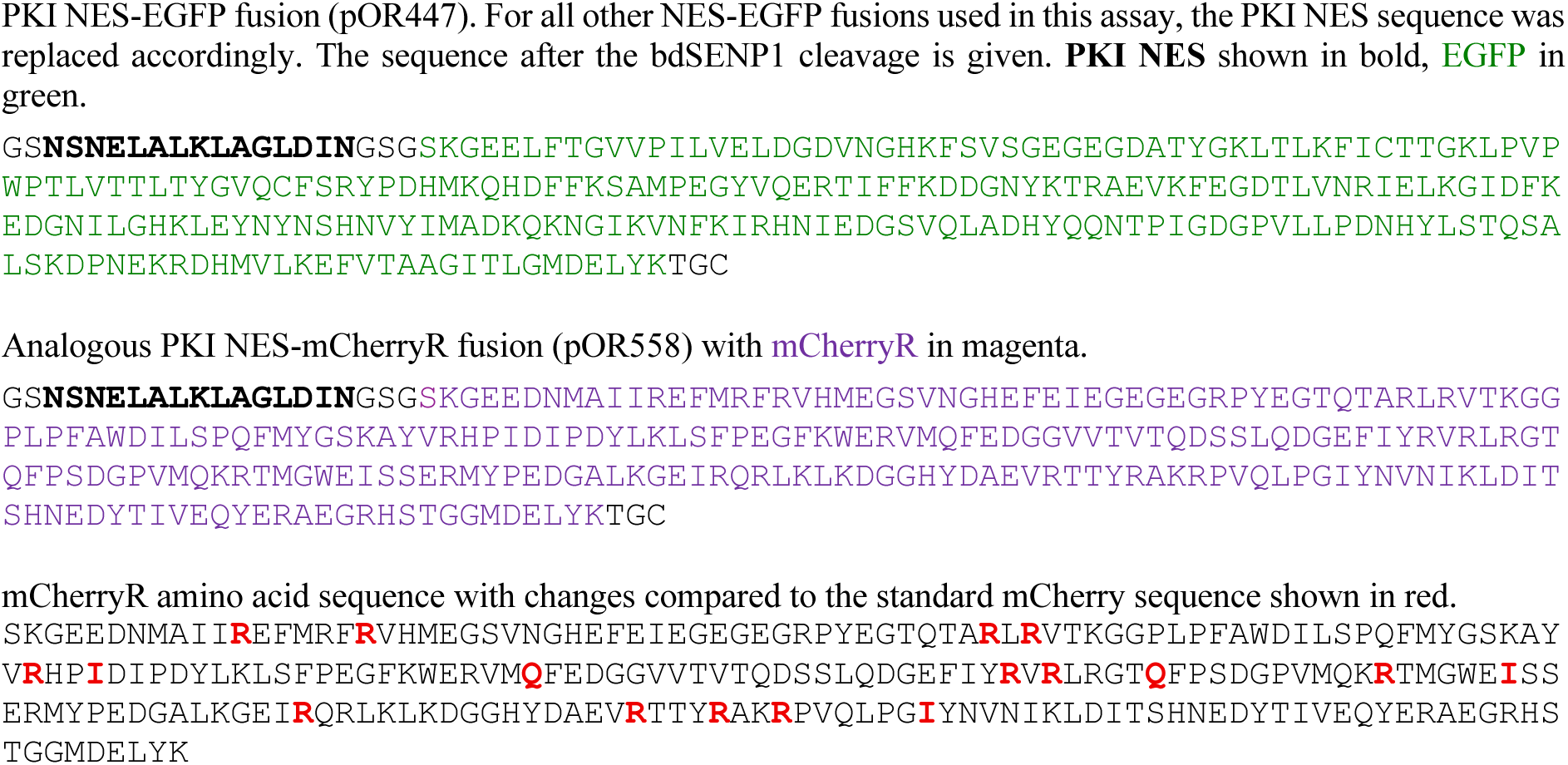

#### Fusion proteins that were transiently expressed in HeLa cells

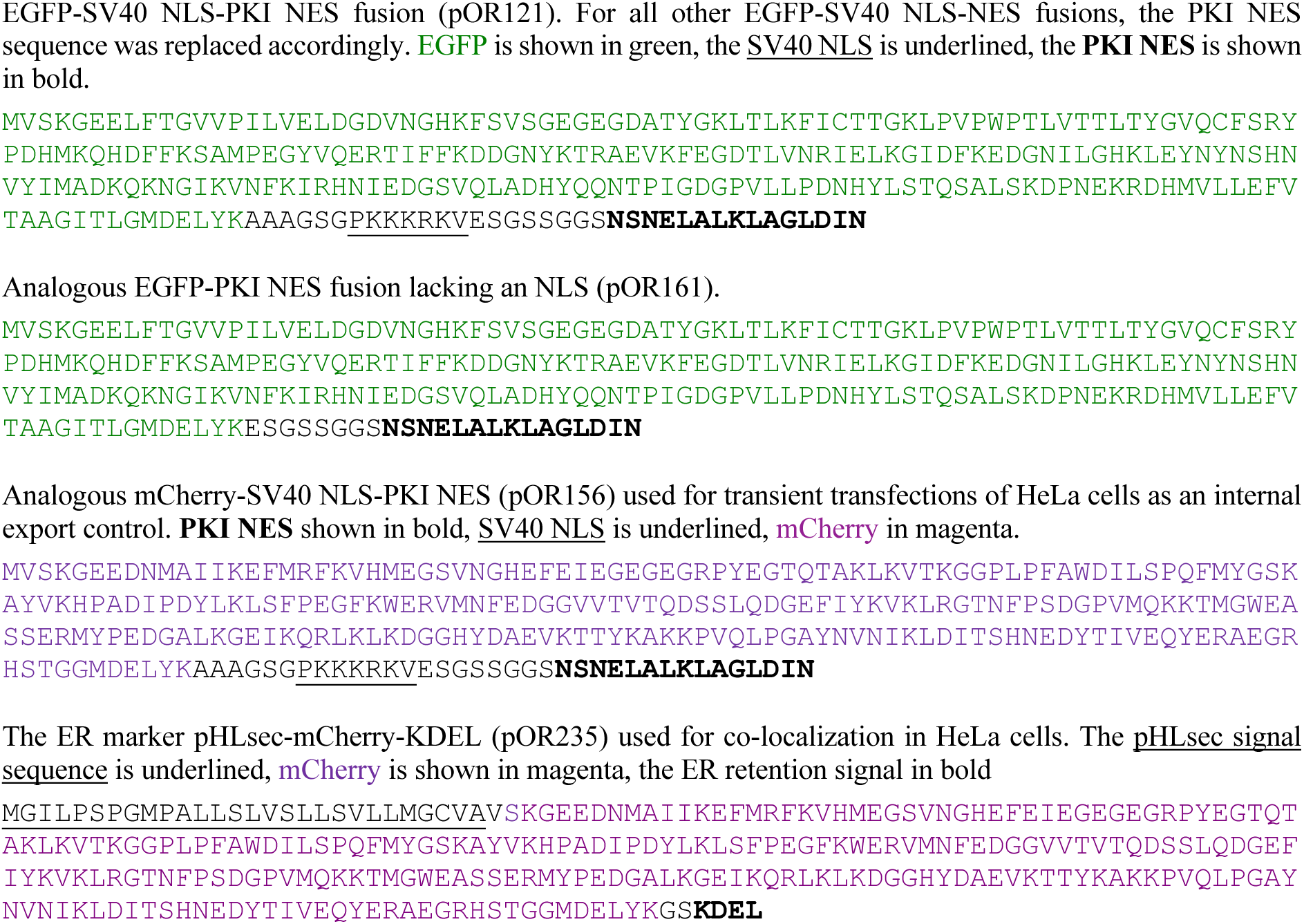

#### NES fusion proteins for fluorescence anisotropy

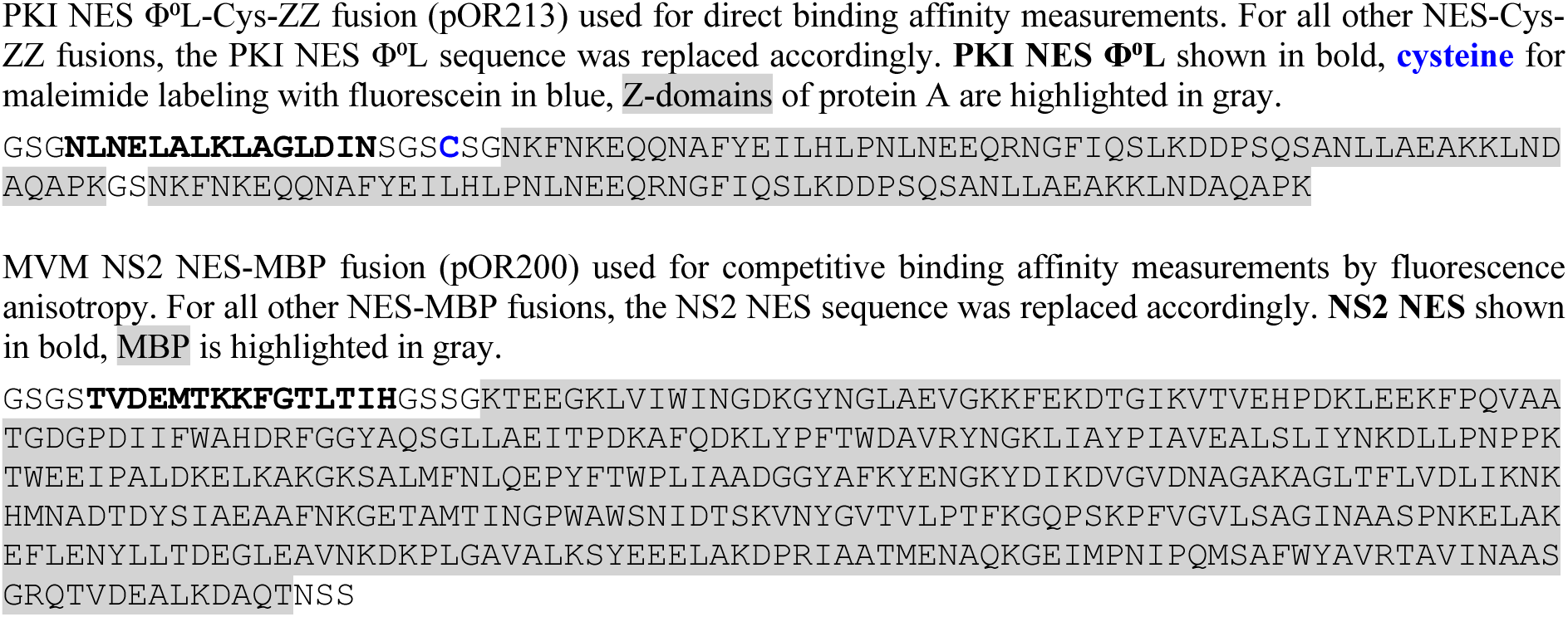

#### Partition coefficient ratio as a relative affinity metric

Note that the following derivation is a simplification of rather complex equilibria and that the simplification is valid only when the Xpo1 is limiting and thus low as compared to the sum of the concentrations of the two NESs. The partition coefficient is defined as a ratio of the mean signal of the fluorescently labeled NES inside the FG phase to that in the adjacent buffer (Figs. 2 and EV2C). Since the signal in the phase originates essentially only from the ternary export complex, while the signal in the surrounding buffer comes mainly from the free NES (provided the exportin concentration is sufficiently low), the partition coefficient *P* is proportional to the ratio of the concentrations of the export complex and the free NES. Considering EGFP-NES1 and mCherryR-NES2 fusions, this yields:

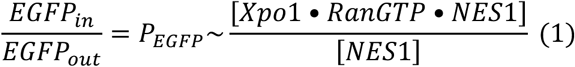

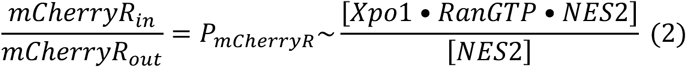

The K_D_s for NES binding to Xpo1 in the presence of a saturating RanGTP concentration can then be written as:

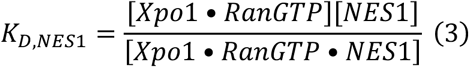

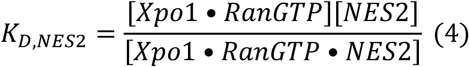

The binding equilibria for the two NESs are coupled because they compete for the same Xpo1·RanGTP complex. Eqs. 3 and 4 can therefore be combined by eliminating the [Xpo1·RanGTP] term. This gives the ratio of the K_D_s:

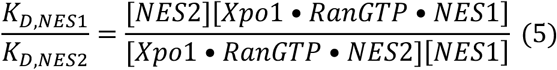

This ratio is thus proportional to the ratio of partition coefficients directly measured by the assay, as defined by the Eqs. 1-2.

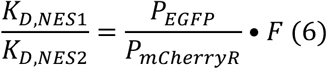

The proportionality factor *F* can be derived by performing the experiment with the same NES being fused to EGFP and mCherryR, which is always included as a control. The proportionality factor is close to 1 in the system with EGFP and mCherryR (Figs. 2C, 3C, 4B. 4D, 8, EV2B).

